# Variation of Human Transfer RNA Demand and Supply

**DOI:** 10.64898/2026.02.26.708130

**Authors:** Marius Külp, Anna-Sophia Wiegand, Halvard Bonig, Michael A. Rieger

## Abstract

Codons function as translation units in open-reading-frames of genes to encode for proteins. Transfer RNAs (tRNAs) connect every codon to its cognate amino acid. The genetic code defines the correspondence between codons and amino acids, yet whether individuals share a common quantitative relationship between tRNA supply and codon demand remains incompletely understood.

Here, we established Marathon-tRNA-sequencing combined with matched mRNA-sequencing to reproducibly quantify tRNA and mRNA abundance from blood cells of ten healthy donors. We apply a precision-biology approach quantitatively integrating transcriptomic codon- and corresponding tRNA abundance across all 61 sense codons to compute the relative tRNA supply per codon.

Thereby, we discover that relative tRNA supply varied approximately 1000-fold across codons, with broadly consistent patterns of highly and lowly supplied codons across donors. The extent of interindividual variability differed between codons and was highest for ACG (threonine). Relative tRNA supply also differed among synonymous codons, and genomic preferences did not consistently identify the codons with the highest relative supply.

In summary, our findings reveal that pronounced inter-codon heterogeneity in relative tRNA supply constitutes a broadly conserved feature of human blood cells, with codon-specific degrees of interindividual variation. By distinguishing shared codon-specific patterns from human variation, this study establishes a reference for investigating codon-dependent gene regulation and synonymous sequence variation.

## Introduction

Transfer RNAs (tRNAs) function as molecular adapters, transporting amino acids to the protein translation machinery thereby linking the coding transcriptome with the proteome of a cell. The genetic code facilitates the correspondence between codons and amino acids, but does not determine how frequently individual codons are used. The codon demand of a cell type is largely determined by the number of transcripts of an expressed set of genes. Neither does the genetic code indicate how abundantly their decoding tRNAs are supplied. A shared genetic code therefore raises a quantitative question: to what extent are transcriptomic codon demand, tRNA supply, and their relationship conserved among individuals? Addressing this question requires distinguishing variation between codons from variation among individuals, because substantial differences between codons themselves may represent a conserved feature of human gene expression.

Studies in bacteria and yeast have provided fundamental insights into how codon usage and tRNA availability influence translation and cellular homeostasis^1–4^. Human studies have linked tRNA expression profiles to distinct cell states^5–10^ and characterized differences in tRNA abundance across tissues^11–14^. However, the extent to which healthy individuals share a common pattern of codon-specific tRNA supply relative to demand remains incompletely characterized. In particular, it is unclear which supply-demand relationships are consistently maintained across individuals, and which exhibit greater interindividual variability.

Recent advances in tRNA-sequencing technologies have expanded opportunities to investigate these relationships^11,12,15–21^. Nevertheless, quantitative tRNA profiling remains challenging because of the stable secondary and tertiary structures of tRNAs, extensive nucleotide modifications, high sequence similarity among tRNA species, and the need to distinguish mature tRNAs from tRNA-derived fragments^22^. Reliable assessment of interindividual variation therefore requires reproducible measurements of both tRNA abundance and transcriptomic codon demand.

Here, we established Marathon-tRNA-sequencing to quantify tRNA abundance in peripheral blood cells from ten healthy donors and combined these measurements with matched transcriptomic profiling via RNA-seq. By incorporating codon-anticodon decoding interactions, we estimated relative tRNA supply across all 61 sense codons. We examined differences between codons, including synonymous codons, quantified their variation among donors, and assessed whether genomically preferred synonymous codons also exhibited the highest relative tRNA supply. Thereby, we establish a reference for distinguishing shared codon-specific patterns of relative tRNA supply from interindividual variation in human blood cells.

## Results

### Marathon-tRNA-sequencing reproducibly quantifies tRNA abundance in blood samples

To investigate interindividual variation in tRNA supply, we established Marathon-tRNA-sequencing using the highly processive reverse transcriptase MarathonRT, which efficiently reverse transcribes human tRNAs as demonstrated by ALL-tRNA-seq^11^. However, in advancement to ALL-tRNA-seq, we facilitated the incorporation of unique molecular identifiers (UMIs) to enable deduplication of amplified molecules preventing PCR amplification biases. We collected peripheral blood (PB) from ten healthy donors (HDs), comprising four female and six male individuals aged 21-29 years. Blood was collected directly into an RNA-preserving cell lysis buffer to minimize RNA degradation during sample processing and storage. A tailored computational alignment pipeline accommodated sequence mismatches potentially introduced during reverse transcription of modified tRNA nucleotides (**Fig. 1A, S1, S2**).

**Figure 1.**
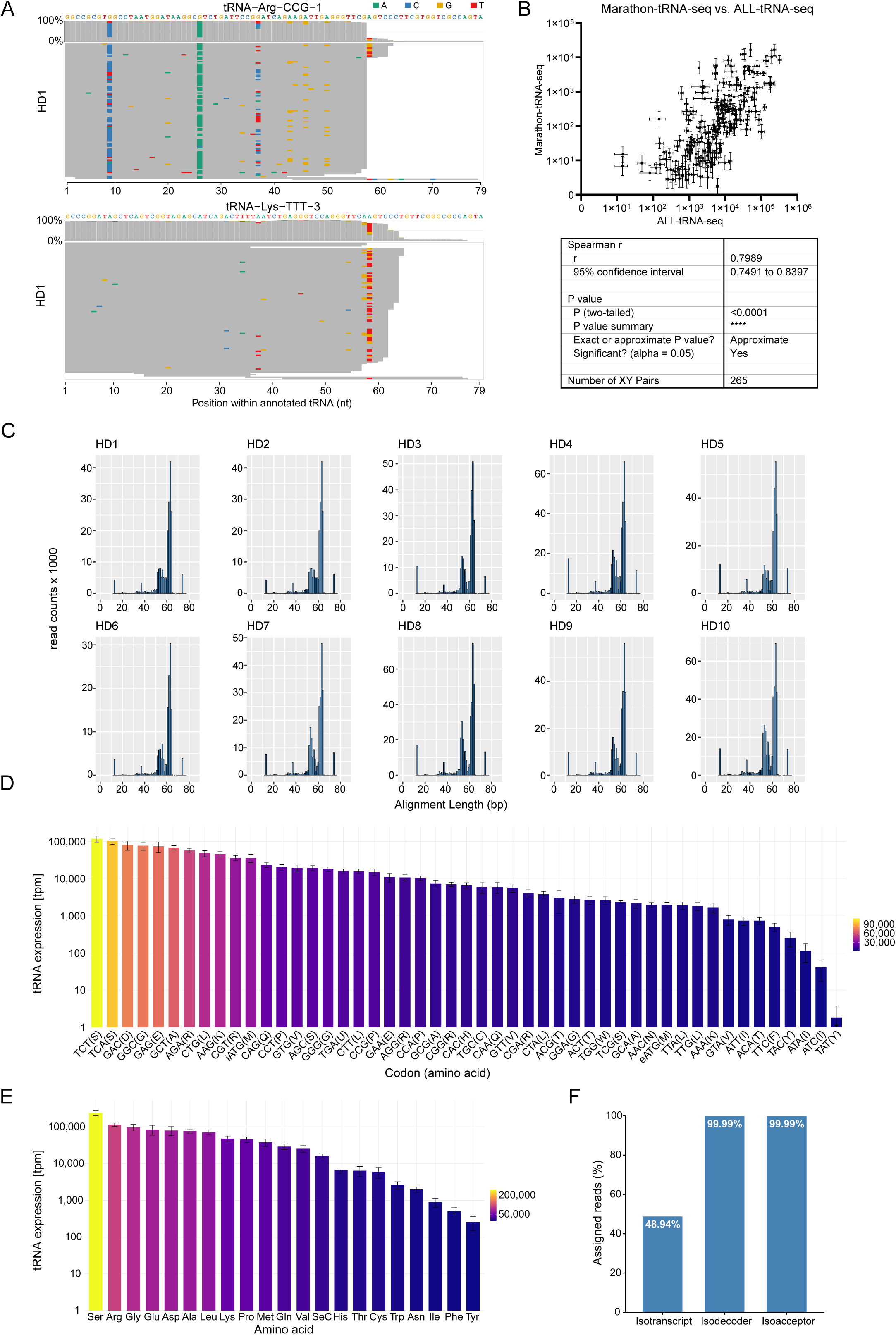
Marathon-tRNA-sequencing reproducibly quantifies tRNA abundance in blood samples. **A** Bam alignment including read coverage and misincorporation signatures of tRNA-Arg-CCG-1 and tRNA-Lys-TTT-3 for HD1. Mismatches are indicated by color (A green, T red, G orange, C blue). Alignment of other donors are shown by **Fig. S1** and **S2**. **B** Comparison and Spearman correlation testing of Marathon-tRNA-seq of peripheral blood cells (n=10 donors) with ALL-tRNA-seq of HEK293T cells (n=3 biological replicates) across tRNA isotranscripts. Correlations across replicates/donors are shown by **Fig. S3A**. **C** Histograms representing the number of tRNA reads against their alignment length before exclusion of reads mapping to less than 30 nt of tRNAs for each donor. **D** Mean tRNA isodecoder expression and corresponding standard deviation per codon normalized to tpm. Corresponding amino acids are shown in one-letter-code. **E** Mean tRNA isoacceptor expression and corresponding standard deviation per amino acid (in three-letter-code) normalized to tpm. **F** Synthetic tRNA-based read assignment analysis. Read assignment specificity of Marathon-tRNA-sequencing of a synthetic tRNA-Lys-CTT-2 is represented as the percentage of read counts assigned to tRNA-Lys-CTT-2 isotranscript, the tRNA-Lys-CTT isodecoder family, or all tRNA-Lys isoacceptors, respectively.

**Figure 2.**
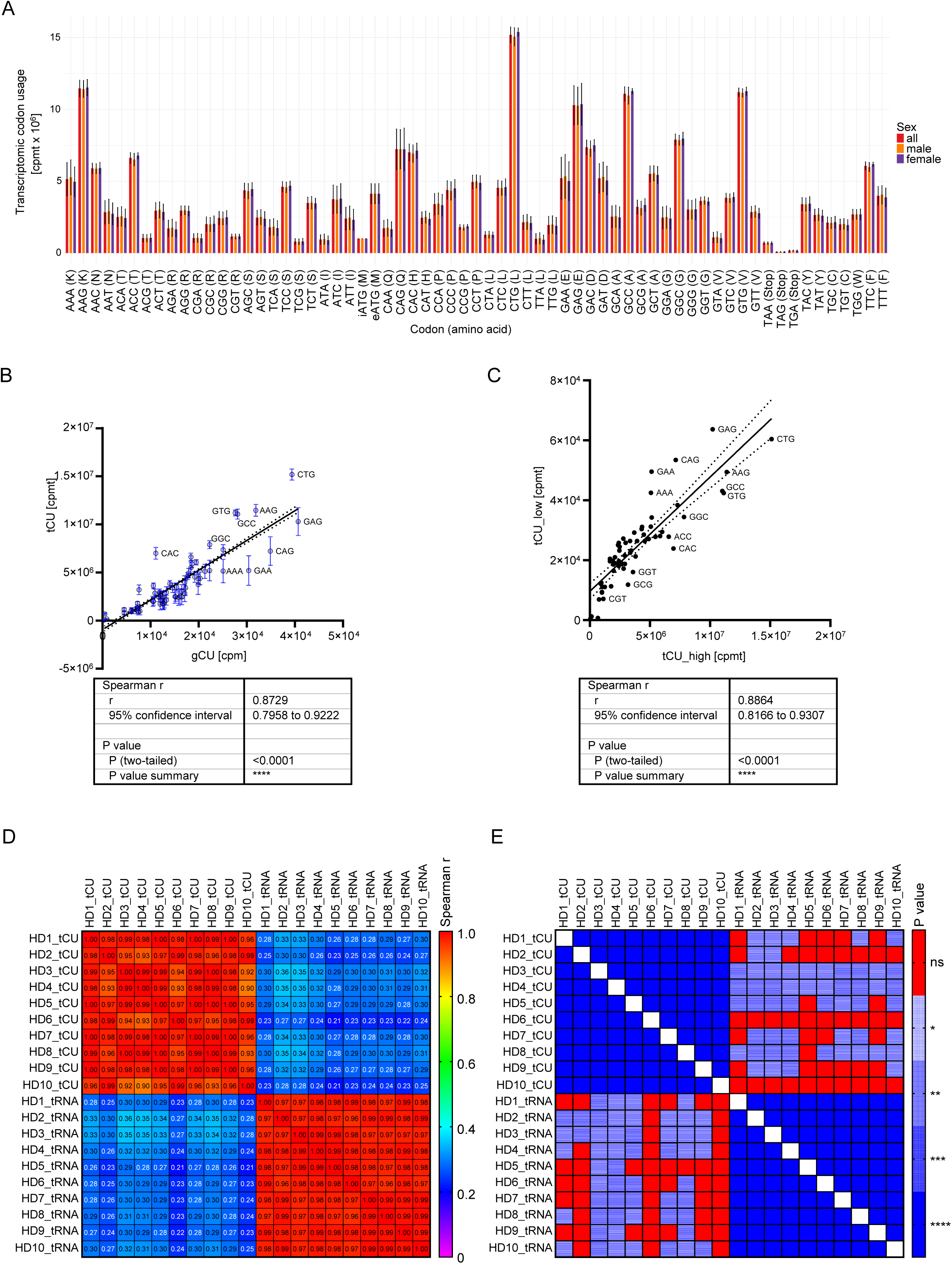
The transcriptomic codon usage (tCU). **A** Codon-wise comparison of tRNA demands. The mean transcriptomic codon usage (tCU) and corresponding standard deviation calculated based on all (red), male (orange) or female (violet) donors is shown. iATG initiator ATG. eATG elongator ATG. **B** Comparison of the tCU in codons per million mRNA transcripts (cpmt) with the genomic codon usage (gCU, in codons per million cpm). Mean and SD of the tCU based on n=10 donors are shown. Each dot represents a codon. **C** Influence of gene expression on the transcriptomic codon usage was inspected using Spearman correlation testing of the tCU calculated based on the top 9,500 highly expressed genes (tCU_high) and the tCU based on the remaining 9,500 genes (tCU_low) across all codons. Each dot represents a codon.

To assess concordance with an established sequencing approach, we compared tRNA abundances measured in blood cells by Marathon-tRNA-seq with published measurements from HEK293T cells obtained using ALL-tRNA-seq^11^. Despite the different biological sources, the tRNA expression profiles correlated strongly (r=0.7989, p<0.0001, **Fig. 1B and S3A**). We further assessed technical reproducibility by sequencing three technical replicates of RNA from a B-cell acute lymphoblastic leukemia (B-ALL) patient-derived long-term culture^23,24^. Strong correlations between replicates demonstrated high technical consistency of Marathon-tRNA-seq (r>0.9868, p<0.0001, **Fig. S3B**).

Alignment length distributions were highly consistent across donors (**Fig. 1C**). Most reads aligned to more than 50 nucleotides, with a modal alignment length of 66-68 nucleotides, indicating that aligned and quantified reads were derived from tRNAs and not tRNA-derived fragments or halves. Reads with alignment lengths below 30 nucleotides were excluded from downstream quantification to limit the contribution of short tRNA-derived fragments.

We quantified tRNA abundance at the levels of individual sequences (isotranscripts), shared anticodons (isodecoders), and amino acid identity (isoacceptors). Abundances differed between anticodon families, with the top 10% most abundant isodecoders (n=5) accounting for 38% of total tRNA (**Fig. 1D**). The serine isodecoders corresponding to the codons TCT and TCA were the most abundant. Consistent with this finding, aggregation by amino acid identity demonstrated serine isoacceptors as the most abundant class (**Fig. 1E**). We detected tRNAs corresponding to all 20 canonical amino acids and selenocysteine.

To assess read assignment specificity, we applied Marathon-tRNA-seq to a synthetic tRNA-Lys-CTT-2. For this target, 99.99% of assigned reads mapped to the expected anticodon family tRNA-Lys-CTT and amino acid class Lys, whereas 48.94% mapped specifically to the input isotranscript tRNA-Lys-CTT-2 (**Fig. 1F**). Read counts assigned to alternative isotranscripts correlated positively with their sequence similarity to the synthetic target (r=0.72, p<0.001, **Fig. S3C,D**).

Notably, principal component analysis (PCA) and unsupervised clustering analysis disclosed no sex-dependent clustering (**Fig. S5**). Moreover, a differential tRNA expression analysis comparing male and female donors yielded no differentially expressed tRNAs, indicating tRNA expression to be sex-independent in blood cells.

Taken together, Marathon-tRNA-seq revealed an unequal distribution of PB tRNA levels across individual sequences, anticodon families and amino acid classes. Clearly, thus, there are highly and lowly expressed tRNAs.

### tRNA demand and supply are imbalanced

Next we aimed to characterize the global tRNA demand to understand the impact of the uncovered differential tRNA supply among codons. The demand for tRNAs is reflected by the expression-weighted frequency of each codon within the coding transcriptome. We refer to this measure as transcriptomic codon usage (tCU), distinguishing it from genomic codon usage (gCU), which solely describes codon frequencies in genomic coding sequences. Expressed as codons per million mRNA transcripts (cpmt), tCU provides an estimate of the relative codon demand for tRNAs (**Fig. 2A**). Among sense codons, tCU varied approximately 19-fold, ranging from 7.97x10^5^ cpmt for TCG (serine) to 1.5x10^7^ cpmt for CTG (leucine). In comparison, the stop codon TAG had the lowest frequency overall, at 7.9x10^4^ cpmt. tCU profiles were similar between female and male donors. Although tCU and gCU correlated across codons (r=0.8729, p<0.0001, **Fig. 2B**), individual codon frequencies strongly differed between the transcriptomic and genomic measures. Disequilibrium was particularly strong for CTG (leucine), GTG (valine), and CAC (histidine)

To examine whether codon usage patterns differ between highly and lowly expressed genes, we calculated the tCU separately for the 9,500 most highly expressed genes (tCU_high) and the 9,500 least expressed genes (tCU_low) in each donor. The resulting profiles were strongly correlated across codons (r=0.8864, p<0.0001, **Fig. 2C**). Thus, although highly expressed genes contributed more strongly to total transcriptomic codon demand, the two gene expression groups exhibited similar relative transcriptomic codon preferences.

We next compared the interindividual variation of tRNA demand and supply. For an initial analysis, we matched codons to tRNA anticodon families based on complementarity (n=47 codon-anticodon pairs) and calculated pairwise Spearman correlations between donor profiles. tCU was highly correlated across the ten donors (**Fig. 2D,E**, upper-left block) and tRNA abundance profiles showed strong interindividual correlations, too (lower-right block). Importantly, despite this interindividual similarity within each dimension, tCU correlated only weakly with tRNA abundance within individual donors (upper-right block). Thus, codons with high transcriptomic demand did not consistently correspond to highly abundant tRNAs. In summary, our analysis identified that for complementary codon-anticodon pairs, tRNA supply and demand are imbalanced in human PB cells.

### The relative tRNA supply differs between codons and is conserved to varying degrees across individuals

Sequence complementarity of codon-anticodon pairs alone does not capture the full decoding capacity of the tRNA pool, because wobble interactions allow individual tRNAs to recognize multiple codons^25,26^. This interaction is influenced by tRNA modifications, including the conversion of adenosine to inosine at position 34 (I34) in eight human tRNA anticodon families: tRNA-Thr-AGT, tRNA-Ala-AGC, tRNA-Pro-AGG, tRNA-Ser-AGA, tRNA-Leu-AAG, tRNA-Ile-AAT, tRNA-Val-AAC, and tRNA-Arg-ACG^27^. We therefore sought to integrate these decoding interactions into a single measure that captures both tRNA supply and transcriptomic codon demand across all 61 sense codons.

To do so, we calculated the normalized translational efficiency (nTE) per codon according to the framework established by Pechmann and Frydman^1^, building on the tRNA adaptation index (tAI)^28^. Effective supply for each codon was estimated by summing the abundances of all recognizing tRNA anticodon families, weighted by interaction-specific decoding efficiencies. These weights accounted for constraints on codon-anticodon pairing, including wobble interactions^28^. For each donor, effective supply was divided by transcriptomic codon usage and rescaled such that the highest codon-specific ratio equaled 1. Thus, nTE represents an estimate of tRNA supply relative to codon demand.

Our model assigned contributions from 47 tRNA anticodon families to all 61 sense codons (**Fig. 3A**). ATG (Met) and TGG (Trp) were the only codons supplied by a single anticodon family that do not recognize additional codons. Each of the eight I34-modified anticodon families contributed to three synonymous codons, representing the broadest decoding repertoire in the model. Median nTE spanned approximately 10^-3^ for TAT (Tyr) to 1 for TCA (Ser).

**Figure 3.**
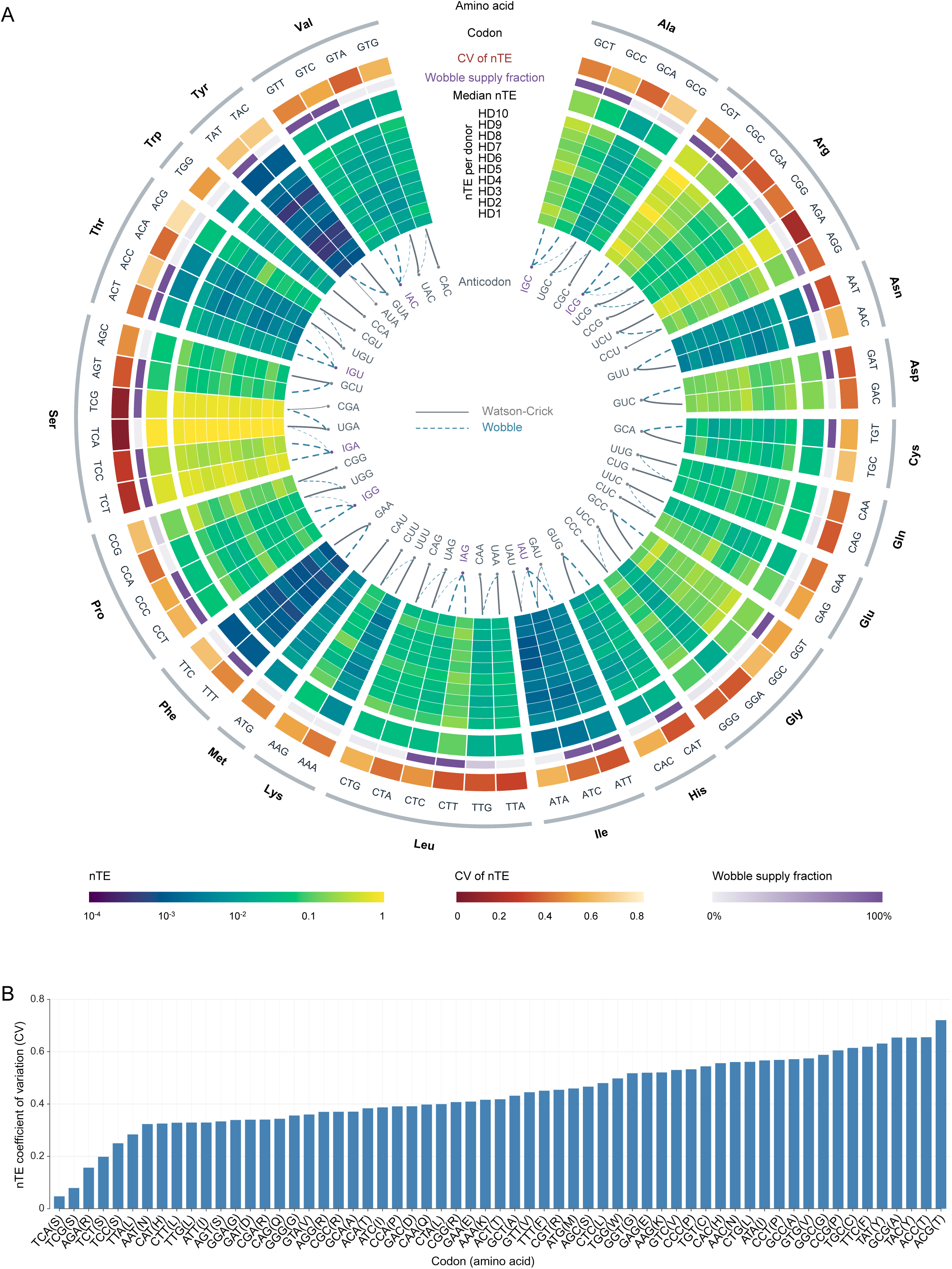
Integration of tRNA supply and demand. **A** Circular assignment of all n=61 anticodon-codon pairs considering Watson-Crick (continuous, grey line) and wobble (dashed, blue line) interactions. Line weight is proportional to the relative decoding contribution. The circular heatmap visualizes donor-specific and median normalized translational efficiency (nTE) values, the fraction of wobble supply relative to total tRNA supply, and the coefficient of variation (CV) of nTE across donors. A-to-I converted anticodons are indicated in violet. **B** The coefficient of variation (CV) across donor nTE values is visualized as a bar diagram with codons in ascending order. Amino acids are indicated in one-letter-code.

The median contribution of wobble interactions to total supply ranged from 0% to 100% across codons. 16 codons had no wobble contribution, whereas 22 received their entire supply through wobble interactions. The remaining 23 codons received Watson-Crick and wobble contributions, with median wobble fractions ranging from 0.01% for TCA (Ser) to 98.67% for TAT (Tyr).

The circular heatmap revealed differences in the interindividual variability of codon-specific nTE values (**Fig 3A**). We quantified this variability using the coefficient of variation (CV) across donors (**Fig. 3A,B**). ACG (Thr) showed the greatest interindividual variation, with a CV above 0.7, whereas TCA (Ser) and TCG (Ser) had the lowest observed CVs, both below 0.1.

Pairwise statistical comparisons across all codons identified substantial differences in relative supply (**Fig. 4**). The five codons with the highest relative supply (TCA (Ser), TCG (Ser), AGA (Arg), TCT (Ser), and CGT (Arg)) had significantly higher nTE values than the 24 lowest-ranked codons, extending from TAT (Tyr) to GTG (Val). In nine donors, TCA (Ser) had the highest nTE, whereas CGT (Arg) ranked highest in the remaining donor.

**Figure 4.**
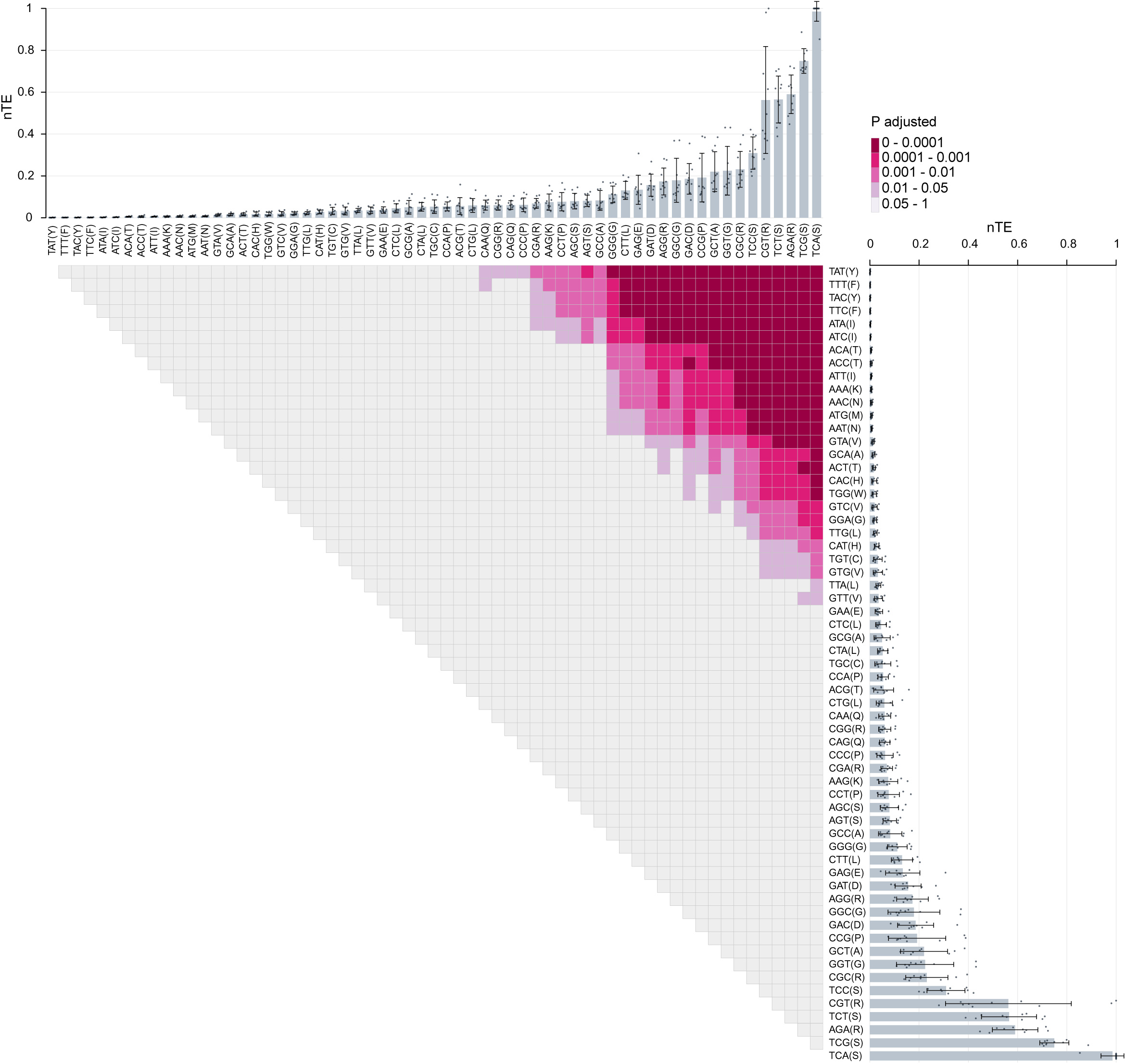
Statistical comparison of relative tRNA supply across codons. Codon-specific nTE is depicted as a bar diagram indicating mean nTE and corresponding standard deviations (error bars). Donor-individual nTE values are indicated as dots. nTE values were compared across n=61 sense codons and n=10 donors, and overall differences assessed using Friedman’s test with a chi-squared approximation. Pairwise comparisons were performed using the repeated measures formulation of Dunn’s test and are visualized as a matrix with color indicating the Bonferroni-adjusted P value. Statistical test results are summarized in **supplementary file S1**.

Taken together, these findings identify two dimensions of variation: relative tRNA supply differs substantially between codons, and the extent of interindividual variation differs from one codon to another.

### Synonymous codons differ in their relative tRNA supply

The circular nTE heatmap (**Fig. 3A**) and comparisons across all codons (**Fig. 4**) indicated that differences in relative supply extended to synonymous codons. We therefore compared nTE values within each amino acid group (**Fig. 5**). Strong statistical evidence for synonymous differences was observed for Ser and Arg (Friedman tests, p=2.1x10^-9^, and p=4.3x10^-9^, respectively). At a nominal significance threshold of p<0.05, differences were detected in 15 of the 18 amino acid groups containing synonymous codons. No significant differences were detected for Asn, Gln or Phe.

**Figure 5.**
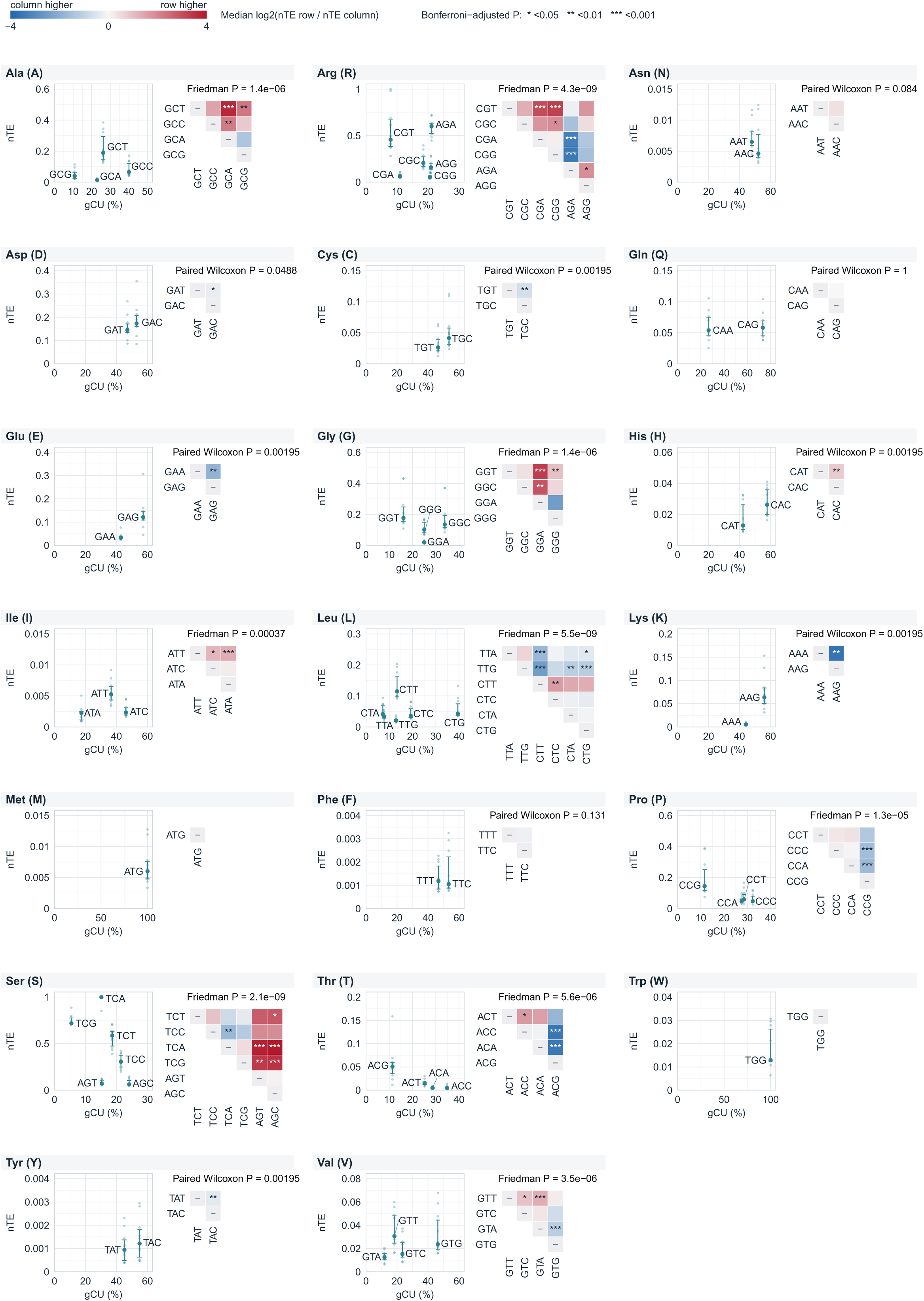
Comparison of relative tRNA supply among synonymous codons and with genomic codon usage. Within each amino acid group, gCU [%] is plotted against median nTE, with individual donor values, and error bars spanning the 25th-75th percentiles. Aside, matrices compare nTE of synonymous codons per amino acid. Cell color represents the median across donors of the log2-transformed nTE ratio between the row and column codons. Two-codon amino acids were assessed using two-sided exact Wilcoxon signed-rank tests, three-or more-codon amino acids were assessed using Friedman tests. Pairwise comparisons were performed using repeated-measures Dunn tests (Bonferroni-corrected). Asterisks indicate Bonferroni-adjusted pairwise significance as * 0.05>p≥0.01, ** 0.01>p≥0.001, *** 0.001>p. Statistical test results are summarized in **supplementary file S2**.

Because synonymous codons also occur at unequal frequencies in genomic coding sequences^29^, we next asked whether genomic codon preference corresponds to higher relative tRNA supply. We compared median nTE values with the genomic fraction of each codon (gCU) within its synonymous family (**Fig. 5**). Among amino acid groups with detectable nTE differences, genomic codon frequencies and median nTE values showed concordant rankings for Asp, Cys, Glu, His, Lys, and Tyr. For Arg, AGA ranked highest in both measures, although the ordering of the remaining synonymous codons differed. In contrast, Ala, Gly, Ile, Leu, Pro, Ser, Thr, and Val showed disagreement between the most frequent genomic codon and the codon with the highest median nTE.

Gln provided a further example of the distinction between genomic frequency and relative supply. CAG accounted for 73.32% of genomic Gln codons, compared with 26.68% for CAA, yet no difference in nTE was detected between them (median nTE_CAG_=0.058 and nTE_CAA_=0.054; paired Wilcoxon test, p=1).

Thus, synonymous codons frequently differ in their demand-normalized tRNA supply, and the most frequent genomic codon does not consistently have the highest relative supply in human blood cells.

## Discussion

Our study identifies a broadly shared, codon-specific organization of tRNA supply relative to demand in human blood cells. This addresses a fundamental question beyond the conservation of codon-amino acid assignments: whether individuals also share the quantitative relationship between the codons expressed by their transcriptomes and the tRNAs available to decode them. By jointly profiling tRNA abundance and transcriptomic codon usage in ten healthy donors, we demonstrate that substantial differences of the relative tRNA supply between codons coexist with considerable similarity among individuals. The central finding is therefore that this unequal distribution of the relative tRNA availability can itself represent a conserved feature of human gene expression. The differences among synonymous codons further demonstrate that this heterogeneity affects codons specifying the same amino acid, too.

With these insights, our study establishes a reference for investigating codon-dependent gene regulation and synonymous sequence variation. Due to the influence of codon composition on translational fidelity, protein folding and mRNA stability, codon usage is regarded as a secondary genetic code^30^. The translation elongation rate per codon directly depends on the supply of the cognate tRNA as the ribosome must pause longer for a scarce tRNA to arrive at the A-site^31–35^. By relating tRNA supply to transcriptomic codon demand, nTE provides a quantitative estimate of codon optimality, consistent with the framework established by Pechmann and Frydman^1^. Historically, the genomic codon usage (gCU) has frequently been used as a surrogate for codon optimality^36^. This is of particular interest as we identified many disproportionalities between nTE and gCU among synonymous codons. As an example, for serine, TCA exhibited the highest nTE despite accounting for only approximately 15% of genomic serine codons. Conversely, AGC was the most frequent genomic serine codon, accounting for approximately 24%, but exhibited one of the lowest nTE values within this amino acid group. Glutamine illustrated a different discrepancy: CAG and CAA showed similar median nTE values despite substantially different genomic frequencies of approximately 73% and 27%, respectively. Thus, gCU does not consistently identify the synonymous codon with the highest tRNA supply relative to demand in blood cells.

However, a higher tRNA supply leading to faster translation does not necessarily maximize the production of correctly folded, functional proteins. Locally slower decoding can facilitate co-translational folding by providing time for nascent polypeptides to acquire appropriate conformations^1,31,37^. This offers a possible functional context for the maintenance of differences in relative tRNA supply across synonymous codons. Accordingly, strategies for interpreting synonymous variants or designing coding sequences should consider cellular tRNA supply and demand together with codon position and protein folding requirements. Our study provides a quantitative foundation for investigating these relationships. Additionally, our resource may guide the translational optimization of expression cassettes for hematopoietic cell and gene therapies, including factor VIII expression in hematopoietic stem cells of hemophilia A patients^38^ or chimeric antigen receptor (CAR) expression in T-cells of cancer patients^39^.

We systematically monitored tRNA demand and supply in healthy donors to integrate both dimensions, which is a long-standing effort in the community. Berg and Kurland presented a co-dependency of tRNA abundance and codon usage on *E. coli* growth in 1997^40^, while Gingold *et al.* later initially proposed^41^ and subsequently confirmed^5^ co-regulation of tRNA demand and supply depending on cell states in cell lines. Moreover, tRNAs were shown to be differentially expressed in human tissues by various studies^12–14^. The normalized translational efficiency (nTE) framework established by Pechmann and Frydman^1^ normalizes codon-specific estimates of tRNA supply derived from the tRNA adaptation index (tAI)^28^ by transcriptomic codon abundance. The original tAI uses tRNA gene copy numbers as proxies for tRNA abundance. However, gene copy numbers do not reflect tRNA abundance in mammalian cells, where individual tRNA genes differ in their transcriptional activity^42–44^. We therefore calculated nTE using tRNA abundances measured by Marathon-tRNA-sequencing, integrating experimentally determined tRNA supply with matched transcriptomic codon demand.

Our analysis of ten healthy donors reveals that the relative tRNA supply follows broadly shared codon-specific patterns, while the degree of interindividual conservation differs across codons. This variation can arise from differences in tRNA abundance, transcriptomic codon demand, or their combined effects. Genomic variation represents one potential source of these differences, particularly because tRNA gene loci experience elevated rates of mutagenesis^45^. The broad conservation of these profiles is consistent with regulatory mechanisms that maintain characteristic relationships between tRNA supply and demand. Many tRNAs are encoded by multiple genes distributed across different genomic locations, such that their abundance reflects the combined transcriptional output of several loci^46^. This organization provides a potential basis for buffering changes in individual gene activity. Other determinants of tRNA gene activity are the sequence-context^47,48^, chaperone binding^49^, and mTORC1 signaling, regulating RNA polymerase III through MAF1^8^. Shared regulatory programs could therefore maintain similar tRNA pools across individuals, while differences in the sensitivity of individual loci to genetic variation could produce unequal variability among decoding tRNA families. Because codons receive distinct combinations and proportions of these tRNA contributions, the resulting stability of supply may itself be codon-specific.

In summary, our study reveals that pronounced differences in relative tRNA supply across codons form a broadly conserved pattern in human blood cells, with codon-specific degrees of interindividual variation. Integrating tRNA abundance and transcriptomic codon demand establishes a physiological reference that genomic codon preferences alone cannot provide.

### Limitations of the study

While previous studies already resolved cell type-specific tRNA patterns^5–10^, our study focuses on their variation among individuals. Our analyses are based on human blood cells derived from healthy young adults and thereby reflect human variation to a limited extent. However, the number of included donors is sufficient to demonstrate a previously unknown dimension of interindividual heterogeneity – and conserved inter-codon heterogeneity.

This research attempt was for now restricted to PB cells because samples are lowly invasive, easily collectable without stress, and available from young and objectively healthy donors. In addition, blood can be easily sampled into an RNA-stabilizing buffer. This is of special interest for tRNA studies on primary human material to prevent degradation-induced tRNA-derived fragments.

Our analyses included Watson-Crick and wobble codon-anticodon pairs, including inosine-dependent decoding. Although other tRNA modifications can influence codon recognition and decoding efficiency^22,50^, too, their quantitative effects remain incompletely characterized across human tRNAs and cellular contexts. Thus, our study addresses a distinct dimension by jointly quantifying tRNA abundance and transcriptomic codon demand and comparing their relationship within a common decoding framework.

## Methods

### Blood sampling and RNA extraction

2 mL residual whole peripheral blood left over after laboratory testing for donor eligibility were collected from young (third decade of life) healthy volunteer donors at the time of eligibility assessment between July and August 2024 at the DRK Blutspendedienst Baden-Württemberg-Hessen in Frankfurt (Main) according to vote #329/10 of the Ethics Committee of Goethe University Medical School. Donors were objectively healthy with no relevant past medical history, no acute complaints, entirely normal physical examination, normal blood counts and extensive normal biochemical and infectious disease laboratory workup^51^. The leukocyte counts and differentials of the healthy donors were highly similar (**Tab. S1**). All donors were Caucasian. Donor sex and age are given in **Table 1**. Samples were collected into EDTA tubes and transferred within one hour into a PAXgene Blood RNA Tube (Qiagen, 762165) to prevent degradation and to stabilize intracellular RNA at room temperature. Samples were frozen below -70 °C on the same day and stored until RNA preparation. Total RNA was extracted and purified using the PAXgene Blood RNA Kit (Qiagen, 762174) according to the manufacturer’s protocol.

**Table 1.**
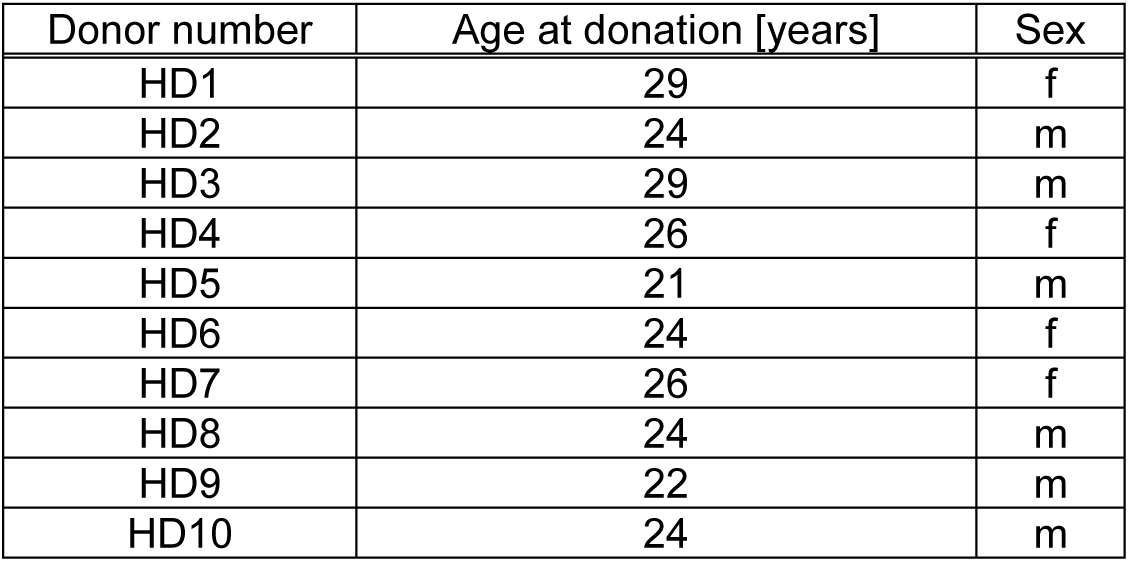
Characteristics of healthy donors. f female. m male.

| Donor number | Age at donation [years] | Sex |
| --- | --- | --- |
| HD1 | 29 | f |
| HD2 | 24 | m |
| HD3 | 29 | m |
| HD4 | 26 | f |
| HD5 | 21 | m |
| HD6 | 24 | f |
| HD7 | 26 | f |
| HD8 | 24 | m |
| HD9 | 22 | m |
| HD10 | 24 | m |

### Marathon-tRNA-sequencing

We developed Marathon-tRNA-sequencing as an integrated experimental and computational workflow for quantitative tRNA profiling. The experimental protocol was established by adapting the TrueQuant smallRNA-Seq Kit (GenXPro GmbH, Germany) used in conjunction with the ultra-processive MarathonRT^11^.

3’ and 5’ adapters containing TrueQuant unique molecular identifiers (UMIs) were sequentially ligated to RNA molecules, followed by reverse transcription using MarathonRT. Libraries were PCR-amplified using the minimally required cycle number and purified using paramagnetic silica beads. Adapter ligation, amplification, and purification followed the manufacturer’s instructions. Library quality and purity were assessed by capillary electrophoresis (Agilent TapeStation system), and libraries were sequenced on an Illumina NextSeq with 1x76 bp. Sequencing data were processed using a dedicated computational pipeline developed alongside the experimental protocol, integrating mismatch-tolerant alignment to tRNA reference sequences, UMI-based deduplication, and quantification of tRNA abundance.

### tRNA sequencing data processing

Processing steps were performed using the public server at usegalaxy.org^52^. Using UMIs, PCR-derived copies were eliminated from raw data and sequencing reads adapter-trimmed and quality-trimmed using Cutadapt (4.6^53^) with the arguments “-e 0.1 -O 3 -q 20 -m 20 -n 8”. Additional sequencing artifacts were removed from the reads with the arguments “-m 20 -u 4 -a ’A{300}X’ -a ’A{10};o=10’”. FastQC (0.12.0^54^) was used to assess the quality of sequencing reads.

An artificial tRNA isotranscript genome was created based on the *Homo sapiens* hg38 high confidence tRNA dataset from GtRNAdb^46^ using the Collapse sequences tool (Galaxy Version 1.0.1) to merge identical tRNA isotranscripts. Collapsed sequences were manually equipped with 5’ (5’-GTTCAGAGTTCTACAGTCCGACGATC-3’) and 3’ (5’-CCAGTACGAGCATCAGTCTAGTCAGTCATCGAAAAAAAAAAAAAAAAAA-3’) dummy adapters. The 3’ dummy adapter introduces the post-transcriptionally added CCA to each tRNA. Two GFF files were manually created. One serves for isotranscript annotation, the other indicates an 11 bp region in each isotranscript starting from tRNA base 32 until 43 (including dummy adapters: base 59 until 70) for filtering purposes. PCR-cleaned and adapter-trimmed FASTQ files are mapped against the artificial tRNA isotranscript genome using Bowtie2^55,56^ with the arguments “-N 1 -L 10 -i S,1,0.5 –local --score-min G,1,8 --np 0”. The resulting mapped reads were filtered using BAM filter to remove reads outside the 11 bp region indicated in the second GFF file with the arguments “-mapped YES -includebed [GFF filtering file]”. This excludes reads not spanning the anticodon loop. Subsequently, featureCounts^57^ was used to count reads assigned to tRNA isotranscripts according to the tRNA annotation GFF file with the arguments “-t tRNA -g transcript_id -M -O --fraction YES --minOverlap 30”. Thereby, only reads overlapping at least 30 bp of a tRNA were counted.

### tRNA transcript normalization

tRNA read counts from featureCounts were normalized to transcripts per million (tpm). The read counts were divided by length of each tRNA isotranscript in kb resulting in reads per kilobase (rpk). The totaled rpk of each sample were divided by 1x10^6^ yielding a scaling factor, to subsequently divide each tRNA’s rpk value by the scaling factor resulting in tpm at isotranscript level (**Fig. S4, Tab. S2**). To inform about tRNA expression at isodecoder level, tpm values of isotranscripts with shared anticodon were summed up and assigned to the corresponding codon (**Fig. 1D, Tab. S3**). The same was performed for isodecoders serving the same amino acid to report tRNA expression at isoacceptor level (**Fig. 1E, Tab. S5**). At each level, tRNA expression was plotted across codons using the R package ggplot2^58^.

### Alignment comparison of tRNAs

BAM-to-SAM^59^ was used to convert the filtered BAM file to SAM format before featureCounts quantification and exclusion of short aligning reads, enabling extraction of the alignment length by summarizing information indicated by arguments “M”, “D”, “N” and “=” using the R script “alignment_length_histogram.R”. Read counts were plotted against the alignment length in a histogram using the R package ggplot2 (**Fig. 1C**). Coverage of the aligned reads was inspected by loading the bam files, artificial tRNA genome file, and the tRNA annotation gff file into the Integrative Genomics Viewer (IGV) for Windows, version 2.19.1^60^ (**Fig. 1A, S1** and **S2**).

### Comparison of Marathon-tRNA-seq and ALL-tRNA-seq

The HEK293T FASTQ files generated by the ALL-tRNA-seq study^11^ (GSE186736) “HEK293T ALL-tRNAseq_1” (GSM5660221), “HEK293T ALL-tRNAseq_2” (GSM5660222), and “HEK293T ALL-tRNAseq_3” (GSM5660223) were retrieved from NCBI GEO and processed as described for the Marathon-tRNA-seq data. tRNA isotranscript read counts of both datasets were assigned in GraphPad Prism version 10.0.0 and tested for Spearman correlation across codons (**Fig. 1B**) and (using a matrix) across donors/samples (**Fig. S3A**).

### Assessment of technical variance

Three technical replicates of the B-cell acute lymphoblastic leukemia (B-ALL) patient-derived long-term culture (PDLTC) CR was profiled using Marathon-tRNA-seq. Spearman correlation testing and plotting was performed with GraphPad Prism version 10.0.0 at isotranscript level (**Fig. S3B**).

### Synthetic tRNA-based read assignment analysis

To assess read assignment specificity, a synthetic human tRNA-Lys-CTT-2 was purchased as a custom RNA oligonucleotide from IDT and processed using the Marathon-tRNA-seq protocol and computational workflow.

Assignment specificity was assessed at three resolution levels: the individual tRNA sequence (isotranscript), anticodon family (isodecoder), and amino acid class (isoacceptor). At each level, specificity was calculated as the percentage of total assigned read counts attributable to tRNA-Lys-CTT-2, the tRNA-Lys-CTT isodecoder family, or all tRNA-Lys isoacceptors, respectively (**Fig. 1F**). To investigate the relationship between sequence similarity and assignment to alternative reference sequences, the distribution of assigned reads across tRNA isotranscripts was examined using a bar diagram (**Fig. S3C**). Percentage nucleotide identity to the synthetic tRNA-Lys-CTT-2 sequence was compared with assigned read counts using Spearman correlation testing in R (**Fig. S3D**).

### Differential tRNA expression analysis

Differential tRNA expression between male and female donors was performed using DESeq2 (1.40.2)^61^. The adjusted p-value (q-value) threshold for considering a difference as significant was set to 0.1.

### Principal component and unsupervised clustering analysis

Principal component analysis at isotranscript level (**Fig. S5A**) was performed using ClustVis^62^. For PCA, unit variance scaling was applied to rows and SVD with imputation used to calculate PCs. The prediction ellipses reflect the area in which a new sample from the same group occurs with probability 0.95. Clustering analysis and heatmap were generated in R using ggplot2 and pheatmap^63^ using a hierarchical clustering algorithm with Euclidean distance and complete linkage. The heatmap was restricted to the top 38 most highly expressed isodecoders for visibility reasons (**Fig. S5B**).

### Library preparation for MACE-sequencing

3’-mRNA Massive Analysis of cDNA Ends (MACE) sequencing libraries were generated using the MACE-Seq Kit (GenXPro GmbH, Germany) according to the manufacturer’s protocol. In brief, mRNA was fragmented, 3’ ends were reversely transcribed and cDNA generated upon template switching and incorporation of TrueQuant UMIs. Subsequently, a PCR was performed with minimal cycle number and the library purified using paramagnetic silica-beads.

### Processing of MACE-sequencing data

Adapter trimming, PCR-copy/artifact removal and quality control were performed exactly as for Marathon-tRNA-sequencing. Processed sequencing reads were mapped to ENSEMBL_dna of hg38 (*Homo sapiens*) using Bowtie2^56^ with the arguments “--sensitive --local”. Quantification of reads mapped to each gene was performed using HTSeq (2.0.2)^64^ with the arguments “-i gene_id -r pos -a 0” and setting strandedness to YES. Raw counts were tpm-normalized as explained in the tRNA-sequencing section.

### Calculation of the transcriptomic codon usage

To calculate the transcriptomic codon usage (tCU), for every gene of the MACE-seq dataset (*G* = 19,462), the count of each of the 64 codons was quantified and weighted with the corresponding gene expression [tpm] of the canonical transcript.

Let *C_i_* reflect the codon *i* and *n*_g,*i*_ be the count of codon *i* in the coding sequence (CDS) of the canonical transcript of a gene *g* with the expression *E*_g_. Then, the weighted codon contribution *S*_g,*i*_ is expressed as:

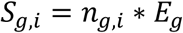

The weighted codon contribution *S*_g,*i*_ is summed for the dataset of *G* genes to calculate the total weighted contribution *R_i_* of codon *i*:

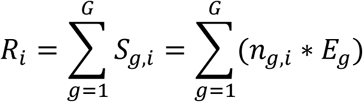

Thereby, the mean transcriptomic codon usage *U_i_* can be calculated in shape of the mean codon expression in codons per million transcripts [cpmt]:

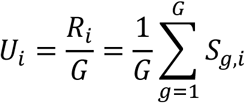

Accordingly, the standard deviation *σ_i_* results as:

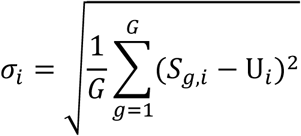

The tCU for every codon was calculated based on the mean expression considering all donors in comparison to exclusively female and male donors, respectively, and subsequently plotted using ggplot2 (**Fig. 2A**).

The tCU was compared with the gCU which was derived from the CAIcal server^29^ by XY assignment and Spearman correlation testing in GraphPad Prism version 10.0.0 (**Fig. 2B**).

For each donor, the tCU was separately calculated based on the top 9,500 expressed genes (tCU_high), and the 9,500 most lowly expressed genes (tCU_low). Spearman correlation testing across the resulting mean tCU of all codons and plotting was performed in GraphPad Prism version 10.0.0 (**Fig. 2C**).

### Integration of tRNA expression and transcriptomic codon usage

To inspect the general correlation of tRNA demand and supply, the Spearman r values and corresponding P values of tRNA expressions and donor-individual tCUs (for anticodon-complementary codons, n=47) were calculated and two matrices plotted using GraphPad Prism version 10.0.0 (**Fig. 2D,E**).

tRNA expression and tCU were integrated for all sense codons (n=61) by calculating the normalized translational efficiency (nTE) independently for each donor according to the framework described by Pechman and Frydman^1^, substituting tRNA gene copy numbers by measured tRNA isodecoder abundances. Our analysis included the 61 sense codons, with ATG supply represented by elongator methionine-tRNA abundance.

A common codon-anticodon decoding matrix was applied across donors. Permitted interactions required Watson-Crick complementarity at the first two codon positions and preservation of amino acid identity. Wobble interactions were incorporated at the third codon position. Complete A34-to-I34 editing was assumed for the eight human ADAT substrate anticodons^27^: tRNA-Ala-AGC, tRNA-Arg-ACG, tRNA-Ile-AAT, tRNA-Leu-AAG, tRNA-Pro-AGG, tRNA-Ser-AGA, tRNA-Thr-AGT, and tRNA-Val-AAC. Reference^28^ tRNA adaptation index penalties were (anticodon:codon):

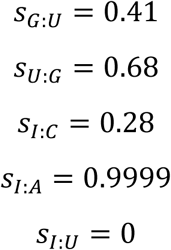

Watson-Crick interactions were assigned *s* = 0.

For each codon, weighted tRNA supply was calculated as

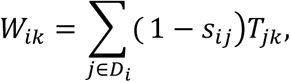

where *T*_jk_ is the measured abundance of anticodon family j in donor *k*, and *D_i_* contains the tRNAs permitted to decode codon *i*. Relative tRNA supply and demand were normalized separately to their respective maxima across all included codons within each donor:

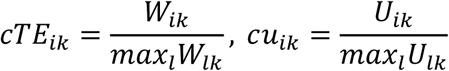

Their ratio was subsequently rescaled to a maximum of one:

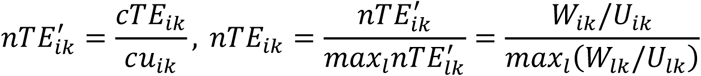

The nTE calculation based on tRNA abundance and tCU was executed using the R script “nTE_framework.R”.

For each codon, the coefficient of variation (CV) of nTE was calculated across 10 donors as the sample standard deviation divided by the arithmetic mean and plotted using the R script “CV_of_nTE.R” (**Fig. 3B**).

A circular heatmap was generated using the R script “plot_nTE_circular.R” to visualize donor-specific nTE values (**Fig. 3A**). The 47 tRNA anticodon families were positioned inside the heatmap and connected to their recognized 61 sense codons according to the described decoding model. Solid lines represented Watson-Crick interactions, whereas dashed lines represented wobble interactions. Inosine-dependent recognition was incorporated by representing the corresponding anticodon position as I34. The median nTE and CV across donors were calculated and visualized by color scales. The wobble supply fraction was calculated separately for each codon and donor as the summed, interaction-weighted contributions of tRNAs recognizing that codon through wobble pairing divided by its total tRNA supply. These fractions were summarized using the median across donors.

Codon-specific nTE was compared across 61 sense codons and 10 donors using a matched design, with donor as the blocking factor. Overall differences were assessed using Friedman’s test with a chi-squared approximation. Pairwise comparisons were performed using the repeated measures formulation of Dunn’s test. Codons were ranked within each donor, and differences in mean ranks were evaluated using two-sided normal-approximation tests. P values were adjusted using the Bonferroni method across all unique codon pairs. Adjusted P values ≤ 0.05 were considered statistically significant. Analyses were implemented in the R script “compare_nTE_codons.R”. Pairwise comparisons were visualized as a matrix with color indicating the adjusted P value and bar diagrams indicating mean nTE values and standard deviations (**Fig. 4**). Statistical test results are summarized in **supplementary file S1**.

### Comparison of relative tRNA supply among synonymous codons and with genomic codon usage

Donor-specific nTE values were compared among synonymous codons within each amino acid group. For amino acids encoded by two codons, differences were assessed using two-sided exact Wilcoxon signed-rank tests. For amino acids with three or more codons, overall differences were assessed using Friedman tests. Pairwise comparisons were performed using repeated-measures Dunn tests, with Bonferroni correction applied separately within each amino acid group.

Pairwise differences were visualized in matrices with cell colors representing the median across donors of the log2-transformed nTE ratio between the row and column codons (**Fig. 4**). Asterisks indicated Bonferroni-adjusted pairwise significance. Statistical test results are summarized in **supplementary file S2**.

Human genomic codon usage (gCU) was obtained from the CAIcal server^29^ and expressed as the percentage contributed by each codon to its synonymous amino acid group. Within each amino acid group, gCU was plotted against median nTE, with individual donor values and error bars spanning the 25th-75th percentiles (**Fig. 4**). Analyses and visualizations were performed in R using the script “synonymous_codon_gCU_comparison.R”.

### Statistical analysis

Statistical analyses were performed using GraphPad Prism version 10.0.0 for Windows, GraphPad Software, Boston, Massachusetts USA, www.graphpad.com, or using R. Tests are specified in the respective methods section and indicated in figure legends. Statistical test results are summarized in **supplementary files S1** and **S2.**

## Resource availability

### Data and code availability

The datasets generated and analyzed during the current study have been deposited in NCBI’s Gene Expression Omnibus and are accessible through GEO Series accession numbers GSE298634 (MACE-seq) and GSE298635 (Marathon-tRNA-seq).

## Acknowledgements

This study was supported by grants from the European Hematology Association (KOG-202409-06466) to MK, the Deutsche Jose Carreras Leukämie-Stiftung (DJCLS15 R/2023) to MAR, the Deutsche Forschungsgemeinschaft DFG (RI 2462/9-1 and RI 2462/10-1) to MAR, and the LOEWE Hessian Funding Program (Hessen State Ministry for Higher Education, Research and the Arts, III L5 − 519/03/03.001 – [0015] and III 5.7 - 519/03/10.001-(0004)) to MAR and HB.

## Author contributions

MK and MAR conceptualized the study and provided funding. MK wrote the manuscript with MAR, performed experimental work and analyzed the data. AW contributed to experimental work. MAR and HB reviewed the manuscript and advised the study. HB collected and provided samples. MAR supervised the study.

