## Supplementary Materials for "Variation of Human Transfer RNA Demand and Supply"

This pdf includes:
Supplementary Figures S1-S5
Supplementary Tables S1-S4

#### Supplementary Figures

**Figure S1. Read coverage of tRNA-Arg-CCG-1 across all donors.**
Bam alignment including read coverage and misincorporation signatures of tRNA-Arg-CCG-1 for all donors (HD1-HD10). Mismatches are indicated by color (A green, T red, G orange, C blue).

**Figure S2. Read coverage of tRNA-Lys-TTT-3 across all donors.**
Bam alignment including read coverage and misincorporation signatures of tRNA-Lys-TTT-3 for all donors (HD1-HD10). Mismatches are indicated by color (A green, T red, G orange, C blue).

**Figure S3. Extended Marathon-tRNA-seq inspection.**
**A** Spearman correlation matrix of Marathon-tRNA-seq of peripheral blood cells (n=10 donors) compared to ALL-tRNA-seq of HEK293T cells (n=3 biological replicates) across donors/replicates for all tRNA isotranscripts. Mean correlations across isotranscripts are shown by **Fig. 1B**. **B** Spearman correlation testing of three technical replicates of PDLTC CR RNA demonstrates low technical variation of Marathon-tRNA-seq. **C** Bar diagram of the synthetic tRNA-based read assignment analysis indicating read counts assigned to the synthetic target tRNA-Lys-CTT-2 and all off-targets. **D** tRNA-assigned reads of the synthetic tRNA-based read assignment analysis as a function of sequence identity to tRNA-Lys-CTT-2, including Spearman correlation testing.

**Figure S4. tRNA isotranscript expression.**
Mean tRNA expression and corresponding standard deviation is shown at isotranscript level.

**Figure S5. tRNA expression as a function of sex.**
**A** tRNA expression at isotranscript level of female (n=4) vs. male (n=6) donors was compared. The prediction ellipses reflect the area in which a new sample from the same group occurs with probability 0.95. The contribution to total variation explained by each principal component (PC) is shown in percent. **B** Unsupervised clustering analysis (hierarchical clustering with Euclidean distance and complete linkage) depending on tRNA isotranscript expression verified sex-independent clustering of samples. The heatmap shows the top 38 most highly expressed tRNAs.

Figure S1

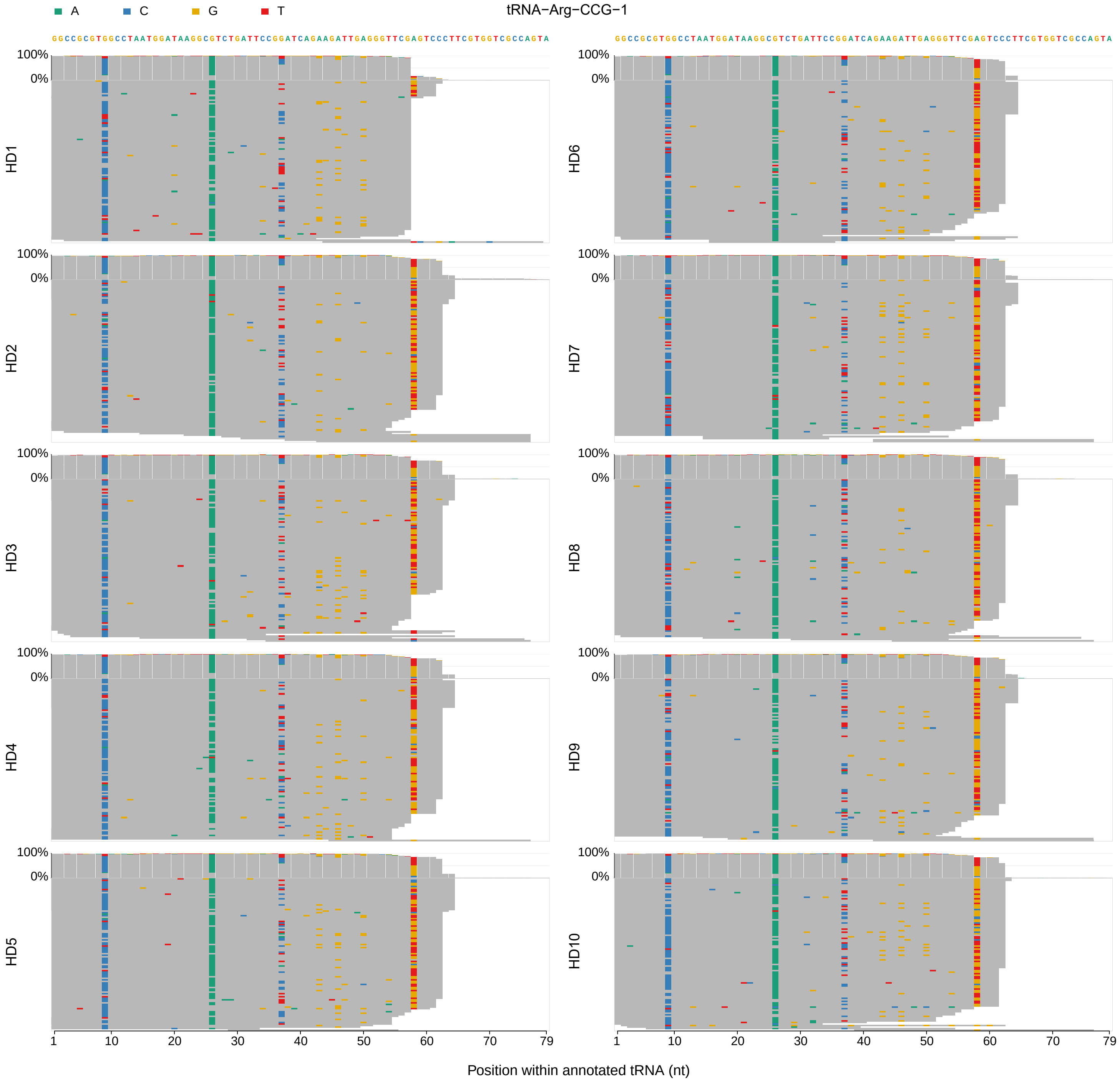
A
C
G
T
tRNA−Arg−CCG−1
GGCCGCGTGGCCTAATGGATAAGGCGTCTGATTCCGGATCAGAAGATTGAGGGTTCGAGTCCCTTCGTGGTCGCCAGTA
GGCCGCGTGGCCTAATGGATAAGGCGTCTGATTCCGGATCAGAAGATTGAGGGTTCGAGTCCCTTCGTGGTCGCCAGTA
100%
0%
HD1
HD2
HD3
HD4
HD5
HD6
HD7
HD8
HD9
HD10
1
10
20
30
40
50
60
70
79
Position within annotated tRNA (nt)

Figure S2

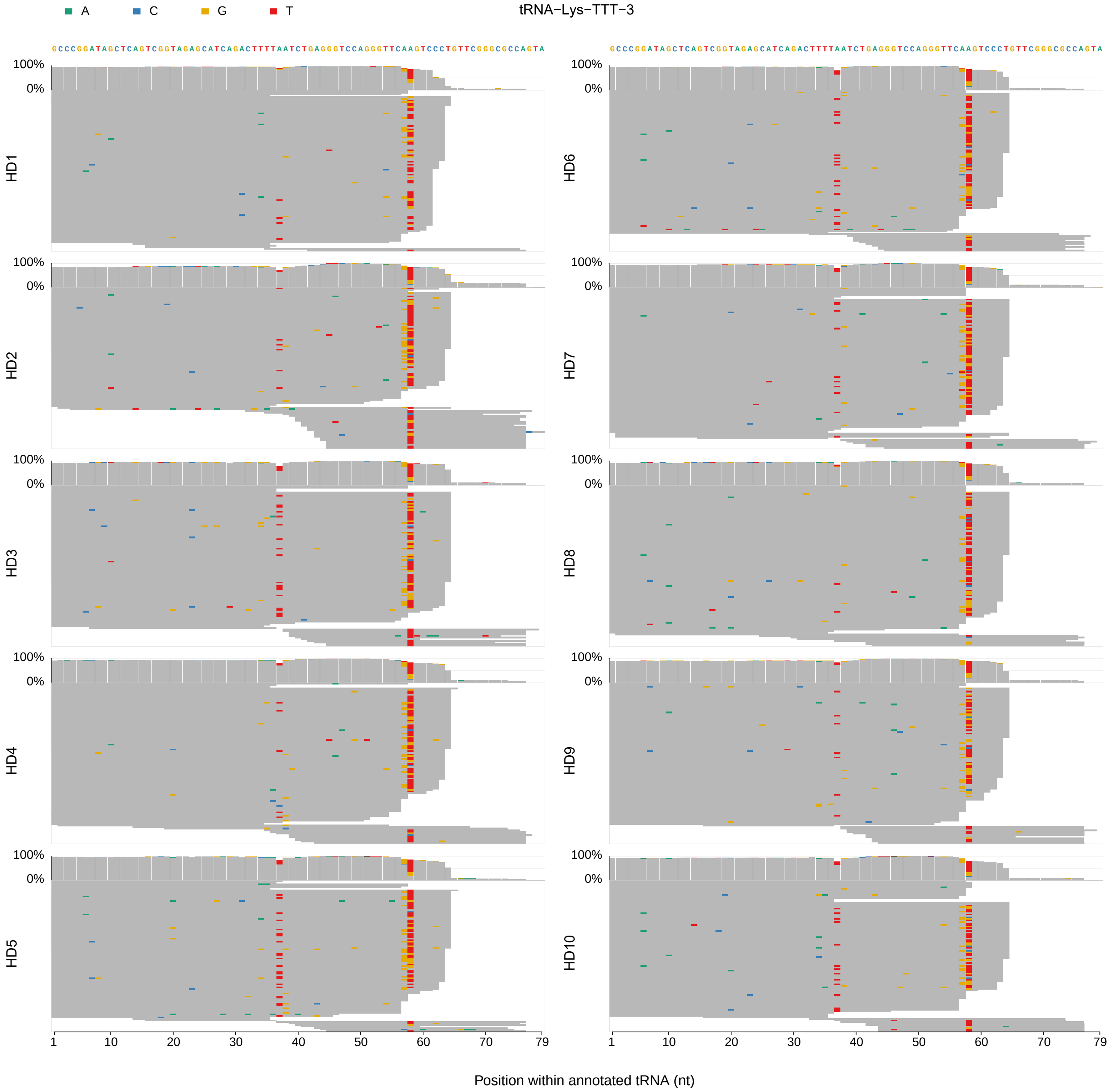
A
C
G
T
tRNA−Lys−TTT−3
GCCCGGATAGCTCAGTCGGTAGAGCATCAGACTTTTAATCTGAGGGTCCAGGGTTCAAGTCCCTGTTCGGGCGCCAGTA
100%
0%
HD1
HD2
HD3
HD4
HD5
HD6
HD7
HD8
HD9
HD10
1
10
20
30
40
50
60
70
79
Position within annotated tRNA (nt)

Figure S3

A

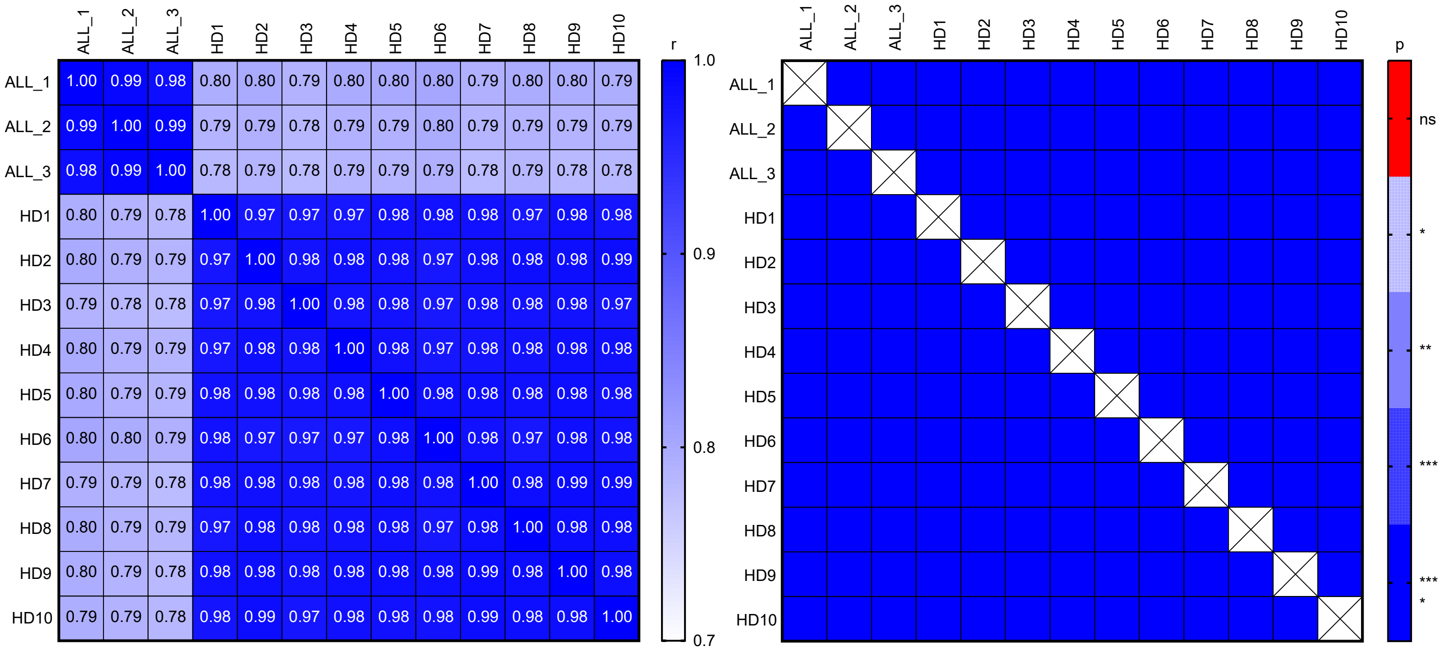

B

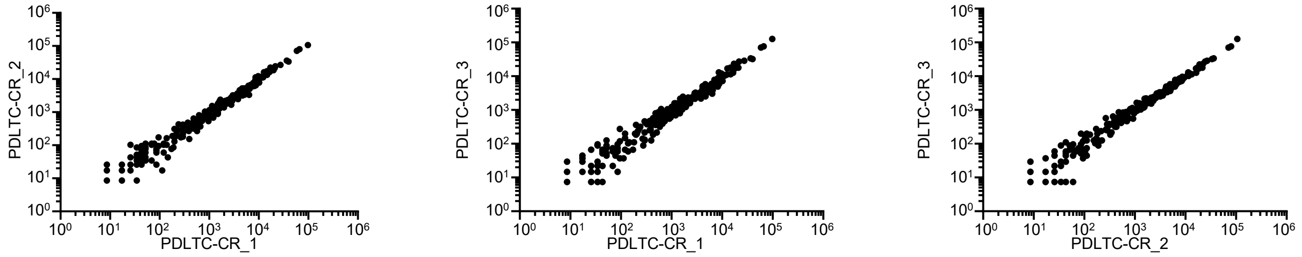

| Spearman r | |
|---|---|
| r | 0.9898 |
| 95% confidence interval | 0.9869 to 0.9920 |
| P value | |
| P (two-tailed) | <0.0001 |
| P value summary | **** |

| Spearman r | |
|---|---|
| r | 0.9868 |
| 95% confidence interval | 0.9831 to 0.9897 |
| P value | |
| P (two-tailed) | <0.0001 |
| P value summary | **** |

| Spearman r | |
|---|---|
| r | 0.9903 |
| 95% confidence interval | 0.9876 to 0.9925 |
| P value | |
| P (two-tailed) | <0.0001 |
| P value summary | **** |

C

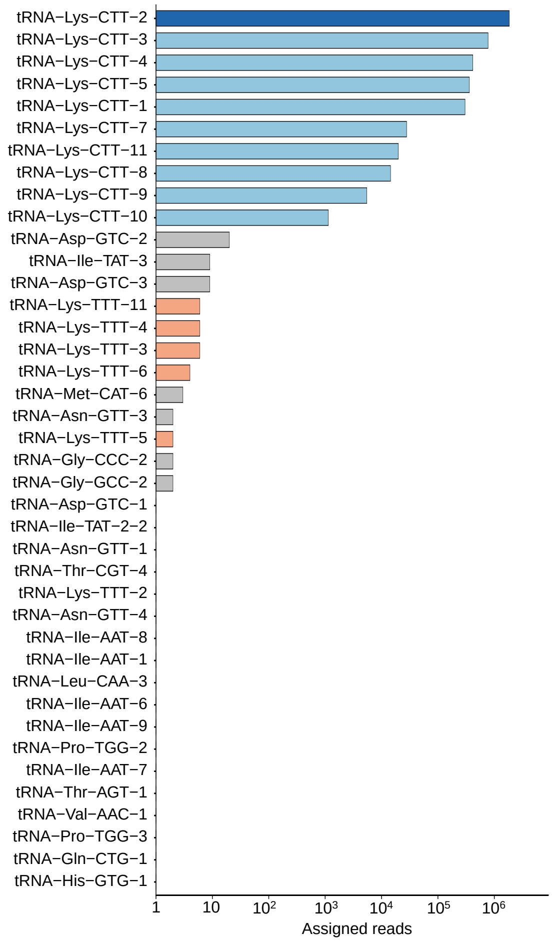

D

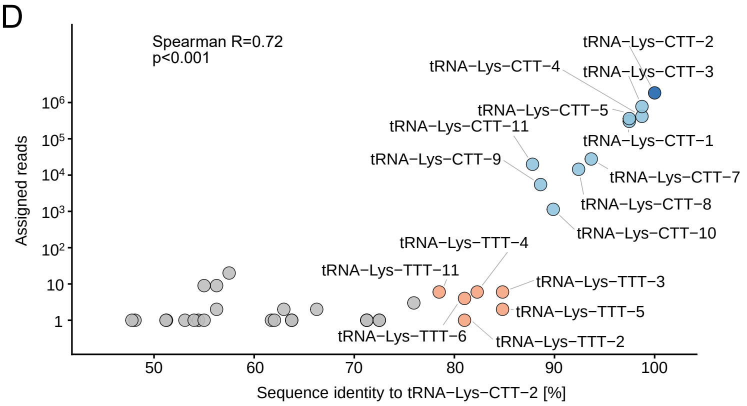

Figure S4

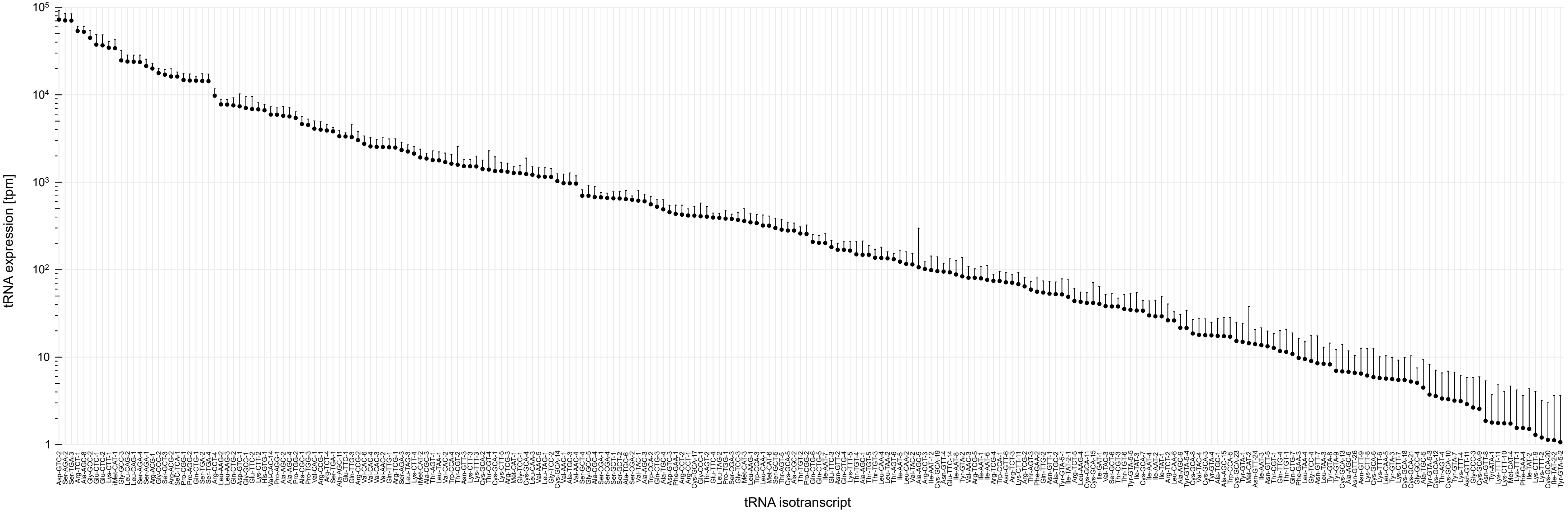

Figure S5

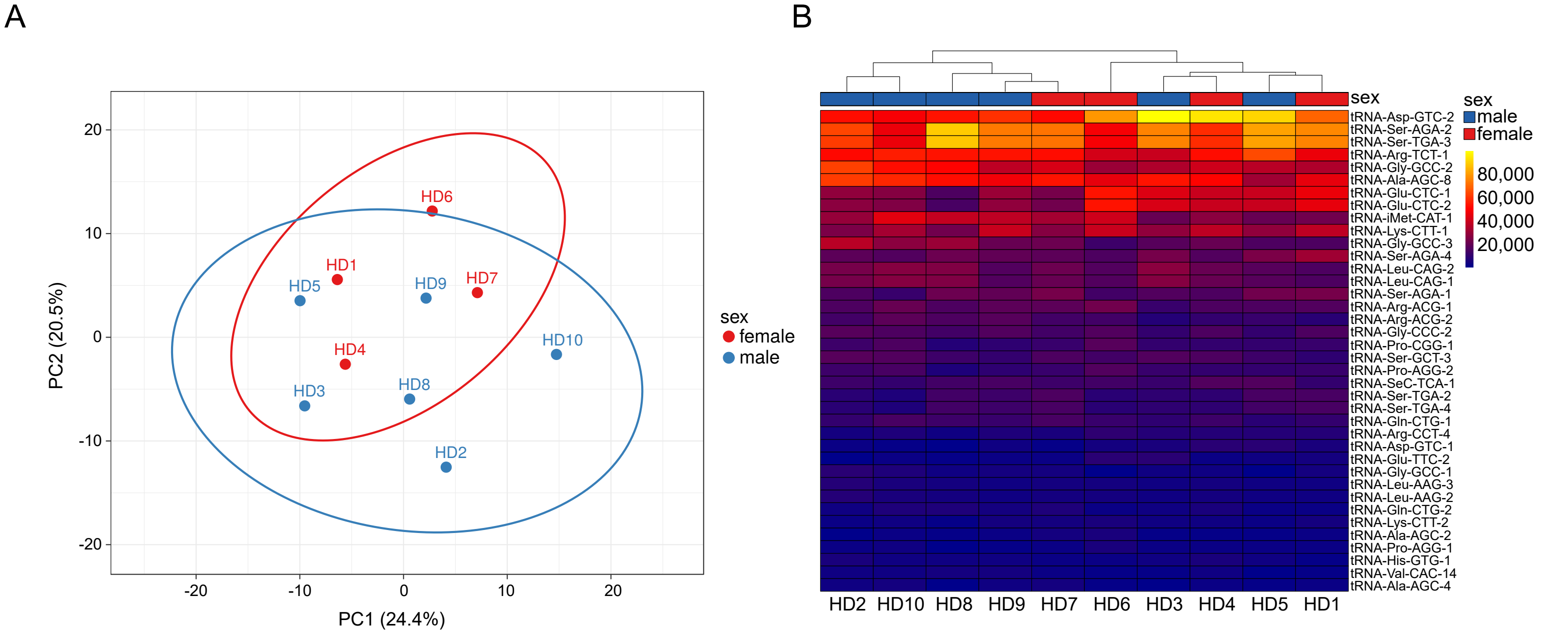
A
B
HD6
HD1
HD5
HD9
HD7
HD4
HD10
HD3
HD8
HD2
PC1 (24.4%)
PC2 (20.5%)
sex
female
male
sex
male
female
80,000
60,000
40,000
20,000
tRNA-Asp-GTC-2
tRNA-Ser-AGA-2
tRNA-Ser-TGA-3
tRNA-Arg-TCT-1
tRNA-Gly-GCC-2
tRNA-Ala-AGC-8
tRNA-Glu-CTC-1
tRNA-Glu-CTC-2
tRNA-iMet-CAT-1
tRNA-Lys-CTT-1
tRNA-Gly-GCC-3
tRNA-Ser-AGA-4
tRNA-Leu-CAG-2
tRNA-Leu-CAG-1
tRNA-Ser-AGA-1
tRNA-Arg-ACG-1
tRNA-Arg-ACG-2
tRNA-Gly-CCC-2
tRNA-Pro-CGG-1
tRNA-Ser-GCT-3
tRNA-Pro-AGG-2
tRNA-SeC-TCA-1
tRNA-Ser-TGA-2
tRNA-Ser-TGA-4
tRNA-Gln-CTG-1
tRNA-Arg-CCT-4
tRNA-Asp-GTC-1
tRNA-Glu-TTC-2
tRNA-Gly-GCC-1
tRNA-Leu-AAG-3
tRNA-Leu-AAG-2
tRNA-Gln-CTG-2
tRNA-Lys-CTT-2
tRNA-Ala-AGC-2
tRNA-Pro-AGG-1
tRNA-His-GTG-1
tRNA-Val-CAC-14
tRNA-Ala-AGC-4
HD2 HD10 HD8 HD9 HD7 HD6 HD3 HD4 HD5 HD1

### Supplementary Tables

#### Table S1. Leukocyte counts and differentials of donors.

| Donor number | WBC/nl | Neutr % | Lymph % | Mono % | Eos % | Basos % | Neutr/nl | Lymph/nl | Mono/nl | Eos/nl | Basos/nl |
|---|---|---|---|---|---|---|---|---|---|---|---|
| HD1 | 5.78 | 57.50 | 27.70 | 11.20 | 3.10 | 0.50 | 3.32 | 1.60 | 0.65 | 0.18 | 0.03 |
| HD2 | 4.09 | 56.90 | 26.70 | 10.00 | 5.40 | 1.00 | 2.33 | 1.09 | 0.41 | 0.22 | 0.04 |
| HD3 | 5.48 | 48.20 | 39.80 | 6.40 | 5.10 | 0.50 | 2.64 | 2.18 | 0.35 | 0.28 | 0.03 |
| HD4 | 4.28 | 54.80 | 33.40 | 8.60 | 2.30 | 0.90 | 2.35 | 1.43 | 0.37 | 0.10 | 0.04 |
| HD5 | 5.72 | 54.30 | 33.60 | 8.70 | 3.10 | 0.30 | 3.11 | 1.92 | 0.50 | 0.18 | 0.02 |
| HD6 | 5.88 | 56.70 | 28.60 | 10.70 | 3.10 | 0.90 | 3.33 | 1.68 | 0.63 | 0.18 | 0.05 |
| HD7 | 6.67 | 46.70 | 44.50 | 6.30 | 2.10 | 0.40 | 3.11 | 2.97 | 0.42 | 0.14 | 0.03 |
| HD8 | 4.63 | 49.80 | 39.30 | 7.80 | 2.20 | 0.90 | 2.31 | 1.82 | 0.36 | 0.10 | 0.04 |
| HD9 | 5.31 | 52.40 | 40.30 | 5.80 | 0.90 | 0.60 | 2.78 | 2.14 | 0.31 | 0.05 | 0.03 |
| HD10 | 4.62 | 57.30 | 27.70 | 10.20 | 3.70 | 1.10 | 2.65 | 1.28 | 0.47 | 0.17 | 0.05 |

#### Table S2. tRNA isotranscript expression [tpm].

| isotranscript | HD1 | HD2 | HD3 | HD4 | HD5 | HD6 | HD7 | HD8 | HD9 | HD10 |
|---|---|---|---|---|---|---|---|---|---|---|
| tRNA-Asp-GTC-2 | 71805.7459 | 54562.4343 | 101817.495 | 96283.1244 | 93710.413 | 81217.4806 | 54476.4802 | 56416.9578 | 62099.6401 | 50902.6698 |
| tRNA-Ser-AGA-2 | 78529.1304 | 65527.0978 | 77465.5439 | 60982.4101 | 83793.8624 | 51063.323 | 74965.3078 | 91292.7453 | 75680.7533 | 49226.4713 |
| tRNA-Ser-TGA-3 | 77779.2144 | 64060.1015 | 77928.8187 | 60867.7873 | 83702.8123 | 50601.8177 | 75046.9366 | 90321.6044 | 76144.7747 | 49574.3615 |
| tRNA-Arg-TCT-1 | 48827.2866 | 53609.6429 | 42535.4856 | 55105.6698 | 66601.2161 | 43813.2244 | 55434.5444 | 56295.9091 | 56384.255 | 57760.3247 |
| tRNA-Ala-AGC-8 | 48537.6639 | 63515.6492 | 56450.8864 | 53243.8243 | 34179.4401 | 48510.1568 | 55602.0983 | 54826.8178 | 49933.2262 | 59826.582 |
| tRNA-Gly-GCC-2 | 37263.0653 | 64755.7905 | 37027.6635 | 42723.313 | 39040.7271 | 33280.4835 | 45475.8314 | 53365.9798 | 40317.0435 | 54555.5175 |
| tRNA-Glu-CTC-1 | 49132.4248 | 31181.2345 | 45739.8026 | 42336.074 | 42548.1378 | 56318.5279 | 26378.9896 | 18575.4766 | 33262.4097 | 29517.9612 |
| tRNA-Glu-CTC-2 | 48382.5089 | 29937.3124 | 45049.18 | 41883.7787 | 42587.7249 | 55656.0446 | 25609.9604 | 18074.7751 | 31560.9981 | 28553.3564 |
| tRNA-Lys-CTT-1 | 40635.1012 | 27086.4999 | 31150.9375 | 40127.2623 | 31424.1829 | 42488.2577 | 29588.2901 | 25128.6143 | 41765.6956 | 35642.9381 |
| tRNA-iMet-CAT-1 | 25228.2072 | 32773.0012 | 23339.6104 | 30275.9001 | 23154.4527 | 43173.0719 | 35371.0458 | 40760.4061 | 40309.4985 | 46717.4446 |
| tRNA-Gly-GCC-3 | 18799.6173 | 39616.4636 | 20903.1284 | 23138.3094 | 18360.4636 | 16108.022 | 24144.9384 | 33040.7989 | 23212.386 | 30045.0676 |
| tRNA-Leu-CAG-2 | 18401.3861 | 26568.514 | 30271.5734 | 23695.9337 | 20474.4109 | 19219.4606 | 24110.5684 | 28325.401 | 19534.1678 | 29517.9612 |
| tRNA-Leu-CAG-1 | 18235.8874 | 26761.3409 | 29353.603 | 23231.2468 | 20126.0451 | 19100.3625 | 23869.9782 | 28688.5471 | 19541.7129 | 29881.6646 |
| tRNA-Ser-AGA-4 | 34397.8692 | 21320.5992 | 26110.6798 | 19724.4097 | 27794.0524 | 19018.4825 | 21928.0721 | 24704.9437 | 22269.2531 | 19165.5905 |
| tRNA-Ser-AGA-1 | 26324.636 | 18299.6453 | 19616.2541 | 18643.2382 | 25078.3823 | 16375.9928 | 26370.3971 | 24680.1838 | 22906.8109 | 15112.1419 |
| tRNA-Arg-ACG-1 | 19637.4544 | 18409.292 | 14859.1087 | 18327.2511 | 19069.0714 | 25464.6688 | 21550.0019 | 18212.3305 | 21148.8113 | 23197.9548 |
| tRNA-Gly-CCC-2 | 18794.4455 | 20466.1116 | 13954.0071 | 18689.7069 | 14069.2298 | 19881.944 | 16879.9756 | 16589.1771 | 19730.3395 | 18406.5572 |
| tRNA-Ser-GCT-3 | 12950.2728 | 20152.2954 | 19517.5937 | 18252.9012 | 15668.5457 | 16316.4437 | 18254.7764 | 15254.8901 | 14090.4049 | 19576.7335 |
| tRNA-Arg-ACG-2 | 12133.123 | 16329.7868 | 10616.7131 | 13404.6679 | 14314.6694 | 17098.0252 | 16669.4592 | 19045.916 | 20616.8843 | 21769.4963 |
| tRNA-SeC-TCA-1 | 13932.9213 | 15724.8399 | 14790.4754 | 19185.3729 | 19239.2957 | 16160.1274 | 15161.4746 | 16223.2799 | 17900.6617 | 13525.5515 |
| tRNA-Pro-CGG-1 | 15060.3812 | 15388.3381 | 14026.93 | 13962.2922 | 13788.1619 | 20261.5693 | 14482.6668 | 10283.6392 | 12226.7744 | 18332.7623 |
| tRNA-Pro-AGG-2 | 15060.3812 | 15645.4406 | 14756.1588 | 13290.0452 | 13518.9701 | 19867.0567 | 14181.9291 | 9782.93763 | 12211.6843 | 17299.6337 |
| tRNA-Gln-CTG-1 | 13591.5803 | 13263.462 | 12195.2789 | 15161.1844 | 11959.2412 | 14299.2192 | 17124.862 | 15582.2719 | 14690.2374 | 17141.5017 |
| tRNA-Ser-TGA-2 | 17377.3629 | 12046.0062 | 12353.9934 | 12506.2733 | 17172.8528 | 11693.9476 | 18486.774 | 16839.5279 | 15716.366 | 9867.43273 |
| tRNA-Ser-TGA-4 | 17811.797 | 11822.932 | 12847.2952 | 12444.315 | 16931.3719 | 11202.6678 | 18082.9263 | 16580.9238 | 15207.0743 | 10141.5281 |
| tRNA-Arg-CCT-4 | 10798.7898 | 7637.45529 | 10346.4695 | 11044.0585 | 10922.0611 | 13138.0124 | 8364.80338 | 6357.80902 | 9548.27708 | 9514.27141 |

| | | | | | | | | | | |
|---|---|---|---|---|---|---|---|---|---|---|
| tRNA-Leu-AAG-2 | 6650.97877 | 10242.5081 | 7094.96703 | 7605.3755 | 6583.32278 | 7495.73852 | 7892.21562 | 8005.7223 | 7118.76683 | 9018.79134 |
| tRNA-Leu-AAG-3 | 6713.04078 | 10287.8791 | 7137.86284 | 7654.9421 | 6765.42311 | 7331.97859 | 7999.62193 | 7529.78074 | 6873.55229 | 9045.14667 |
| tRNA-Gln-CTG-2 | 5714.87678 | 6953.10905 | 6721.77348 | 8389.14739 | 5744.07778 | 5709.26658 | 9683.75286 | 9755.42656 | 7529.97276 | 9498.45821 |
| tRNA-Asp-GTC-1 | 8673.16594 | 5331.09503 | 8652.08495 | 12122.1321 | 11377.3119 | 7845.58928 | 4502.47249 | 3708.49267 | 6530.25193 | 4986.42701 |
| tRNA-Gly-GCC-1 | 7266.42704 | 11683.038 | 6417.21323 | 6428.1687 | 3839.94173 | 3312.41672 | 7973.84442 | 6679.68857 | 7333.80113 | 9677.67441 |
| tRNA-Glu-TTC-2 | 7390.55106 | 3739.32836 | 11200.0961 | 6195.82525 | 6995.02787 | 11902.3693 | 5503.4993 | 4423.78057 | 5677.65982 | 5724.37604 |
| tRNA-Lys-CTT-2 | 8088.74867 | 5652.4731 | 6412.92365 | 7035.35957 | 5292.78566 | 8798.37431 | 7991.02943 | 4869.45996 | 7488.47492 | 6551.93316 |
| tRNA-His-GTG-1 | 4685.68178 | 8253.745 | 7142.15243 | 8274.52463 | 7082.11933 | 5835.80834 | 5847.19949 | 6522.87546 | 6526.4794 | 6309.4642 |
| tRNA-Val-CAC-14 | 4571.90142 | 6862.36701 | 4448.29553 | 6434.36453 | 3839.94173 | 5128.6632 | 6057.71585 | 7722.35825 | 7420.56935 | 6963.07619 |
| tRNA-Pro-AGG-1 | 6144.13902 | 6079.71688 | 5756.61775 | 5405.85753 | 5348.2075 | 8314.53816 | 5714.01566 | 3997.35894 | 5198.54833 | 7216.08729 |
| tRNA-Ala-AGC-2 | 6263.09121 | 3584.3107 | 4714.24956 | 4560.12739 | 5272.99215 | 8478.29809 | 8214.43455 | 4748.41124 | 5681.43235 | 6066.99523 |
| tRNA-Ala-AGC-4 | 5399.3949 | 6385.97128 | 4049.3645 | 5960.38389 | 4829.61743 | 4577.83435 | 7419.62786 | 3345.3465 | 6877.32482 | 7706.29629 |
| tRNA-Pro-TGG-2 | 6713.04078 | 5047.52615 | 4422.55805 | 4724.31676 | 5910.3433 | 7197.9932 | 5413.278 | 4418.27836 | 4753.38962 | 5808.71307 |
| tRNA-Ala-CGC-1 | 3584.08109 | 3879.22234 | 3714.77718 | 5353.19302 | 3341.14517 | 4801.14334 | 6715.04247 | 5386.66813 | 4693.02912 | 4833.56614 |
| tRNA-Pro-TGG-3 | 3966.79682 | 5104.23992 | 3311.55656 | 4368.0568 | 4176.43147 | 5597.61208 | 4854.76519 | 3672.72827 | 4700.57418 | 5455.55175 |
| tRNA-Val-CAC-1 | 3134.13152 | 5459.64626 | 4036.49576 | 4179.08413 | 2735.46364 | 3103.995 | 3883.81215 | 5254.61498 | 4515.72014 | 4754.50017 |
| tRNA-Arg-CCG-1 | 3211.70903 | 3784.69938 | 3002.70673 | 2874.86292 | 3733.05675 | 4049.3364 | 5516.38805 | 4597.10033 | 3866.84473 | 5365.94365 |
| tRNA-Arg-TCT-4 | 3372.03589 | 4030.45908 | 2951.23175 | 4135.71336 | 5170.06587 | 2806.24967 | 4300.54863 | 3986.35451 | 4183.73737 | 4127.2435 |
| tRNA-Ser-TGA-1 | 3609.94027 | 2960.45915 | 4143.73528 | 3556.4037 | 4188.30758 | 3759.03471 | 4051.36599 | 3620.45723 | 4085.65155 | 4274.8333 |
| tRNA-Ala-AGC-11 | 2870.36798 | 4234.62868 | 3809.14796 | 2980.19195 | 2727.54624 | 2888.12964 | 3626.03701 | 3452.63969 | 3067.06806 | 3969.11156 |
| tRNA-Glu-TTC-1 | 3103.10051 | 2971.80191 | 3345.87321 | 3364.33311 | 3194.67317 | 3766.47834 | 2835.52657 | 3887.31464 | 3365.09805 | 3563.2396 |
| tRNA-Gln-TTG-3 | 3408.23873 | 2075.72423 | 1213.95143 | 2143.75554 | 2181.24525 | 4533.17255 | 5185.57662 | 3334.34207 | 4640.21368 | 4100.88817 |
| tRNA-Arg-CCG-2 | 3051.38217 | 2279.89383 | 2921.20469 | 3980.81773 | 3246.13631 | 4548.05982 | 2710.93525 | 1936.77955 | 3055.75047 | 2624.99012 |
| tRNA-Val-CAC-6 | 1784.2828 | 3814.94673 | 3105.65667 | 2633.22573 | 2054.56676 | 2166.09723 | 2741.00902 | 3158.2712 | 2663.4072 | 3384.0234 |
| tRNA-Val-CAC-4 | 1779.11096 | 3792.26122 | 2762.49019 | 2434.95932 | 1623.06815 | 2069.33 | 2350.05005 | 3246.30664 | 2478.55316 | 3178.45189 |
| tRNA-Val-CAC-3 | 1867.03214 | 3149.50508 | 2286.34669 | 2887.25457 | 1555.77021 | 2099.10453 | 2762.49028 | 3100.49795 | 2674.72479 | 3015.04889 |
| tRNA-Val-AAC-2 | 1887.71948 | 3138.16232 | 1750.14906 | 2623.93199 | 1294.49582 | 1987.45003 | 2921.45162 | 2968.4448 | 3565.04221 | 3157.36763 |
| tRNA-Gln-TTG-1 | 2591.08893 | 1909.36382 | 1359.79719 | 2239.79083 | 2137.69952 | 3327.30399 | 3114.78297 | 2599.79642 | 2621.90935 | 3199.53615 |
| tRNA-Arg-TCG-1 | 2058.39001 | 1750.56525 | 1600.01373 | 2422.56767 | 2185.20395 | 3029.55867 | 3385.44687 | 2170.62368 | 3104.79338 | 3231.16253 |
| tRNA-Ser-AGA-3 | 2834.16514 | 2045.47689 | 2243.45088 | 1750.32063 | 3008.61414 | 1578.05021 | 2882.78535 | 2709.84071 | 2587.95657 | 1713.09596 |
| tRNA-Leu-TAG-3 | 1536.03475 | 2514.31078 | 1908.86356 | 2323.43447 | 2030.81454 | 2091.6609 | 2758.19403 | 2495.25434 | 1916.44597 | 2925.44079 |
| tRNA-Lys-CTT-4 | 2497.99591 | 1674.94688 | 1793.04487 | 2317.23864 | 1658.69648 | 2761.58788 | 2246.93999 | 1507.60681 | 2429.51025 | 2398.33434 |
| tRNA-iMet-CAT-2 | 1287.78671 | 2272.33199 | 1308.32222 | 1573.73961 | 1630.98556 | 2106.54816 | 1808.72225 | 2233.89914 | 2410.64759 | 2598.63479 |
| tRNA-Ala-CGC-3 | 1603.2686 | 1663.60412 | 1685.80535 | 1855.64966 | 1456.80263 | 2017.22457 | 2324.27254 | 1934.02845 | 1893.81078 | 2240.20241 |
| tRNA-Thr-AGT-1 | 1086.08518 | 1807.27902 | 2629.51318 | 1477.70432 | 1543.8941 | 1726.92288 | 1821.61101 | 1535.11788 | 1667.4589 | 2640.80331 |
| tRNA-Leu-TAA-1 | 1691.18978 | 1398.93983 | 995.1828 | 1425.03981 | 1528.05929 | 2195.87176 | 2135.23743 | 1994.55281 | 2067.34723 | 2382.52115 |
| tRNA-Val-CAC-2 | 1613.61227 | 2707.13762 | 1848.80943 | 1257.75253 | 997.593109 | 1533.38842 | 1757.16722 | 1625.90443 | 1618.41599 | 2034.63089 |
| tRNA-Trp-CCA-4 | 1432.59807 | 2306.36026 | 887.943275 | 1446.7252 | 1187.61084 | 1764.14104 | 1439.24455 | 1840.4908 | 2195.6133 | 1818.51725 |
| tRNA-Thr-CGT-2 | 2037.70267 | 574.699606 | 2131.92178 | 1954.78287 | 1900.17735 | 3669.71111 | 777.621681 | 308.12402 | 1573.14561 | 932.978415 |
| tRNA-Asn-GTT-3 | 1122.28802 | 1489.68187 | 1690.09493 | 1499.38971 | 1393.46339 | 1875.79554 | 1499.39208 | 1282.01601 | 1358.11132 | 2066.25728 |
| tRNA-Lys-CTT-3 | 1696.36161 | 1285.51228 | 1398.40342 | 1443.62728 | 1076.76716 | 1860.90827 | 1542.3546 | 1232.49608 | 1697.63915 | 2003.00451 |
| tRNA-Lys-TTT-3 | 2389.3874 | 1319.54054 | 776.414168 | 1304.22122 | 1076.76716 | 1838.57737 | 1516.57709 | 1331.53594 | 2074.89229 | 1539.15083 |

| | | | | | | | | | | |
|---|---|---|---|---|---|---|---|---|---|---|
| tRNA-Cys-GCA-2 | 1277.44305 | 1924.4875 | 776.414168 | 1127.6402 | 1187.61084 | 1525.94478 | 1782.94474 | 1213.23833 | 1535.4203 | 1871.22789 |
| tRNA-Thr-CGT-4 | 1970.46883 | 597.385117 | 1651.4887 | 1784.39767 | 1947.68178 | 3178.43133 | 455.402752 | 346.639522 | 1365.65638 | 653.611997 |
| tRNA-Cys-GCA-1 | 992.992165 | 2703.3567 | 737.807938 | 972.744565 | 1021.34533 | 990.003201 | 1370.50451 | 1067.42964 | 1501.46751 | 2076.79941 |
| tRNA-Lys-CTT-5 | 1613.61227 | 1156.96105 | 1089.55358 | 1205.08801 | 989.675703 | 1652.48655 | 1486.50332 | 951.883133 | 1282.66069 | 2003.00451 |
| tRNA-Arg-TCG-3 | 1148.14719 | 1062.43809 | 819.309978 | 1180.30471 | 1041.13884 | 1466.39572 | 1843.09227 | 1378.30477 | 1535.4203 | 1744.72235 |
| tRNA-Met-CAT-1 | 1060.22601 | 1194.77023 | 900.812018 | 1149.32558 | 1104.47808 | 1503.61388 | 1374.80076 | 1370.05145 | 1410.92676 | 1692.0117 |
| tRNA-Gly-TCC-1 | 1179.1782 | 1716.53698 | 870.78495 | 988.234128 | 981.758297 | 1473.83935 | 1391.98577 | 1182.97615 | 1410.92676 | 1528.6087 |
| tRNA-Cys-GCA-4 | 972.304828 | 2620.17649 | 712.070452 | 808.555196 | 791.740562 | 707.145143 | 1177.17315 | 990.398635 | 1584.46321 | 2050.44409 |
| tRNA-Leu-CAA-3 | 1401.56707 | 1028.40982 | 707.780871 | 923.177963 | 1132.189 | 1518.50115 | 1611.09464 | 1108.69625 | 1339.24866 | 1417.91635 |
| tRNA-Val-AAC-5 | 1065.39784 | 1395.15891 | 759.255843 | 1081.17151 | 629.433747 | 1146.3195 | 1194.35816 | 1543.37121 | 1512.78511 | 1317.76612 |
| tRNA-Leu-TAG-1 | 806.806134 | 1315.75962 | 1179.63479 | 957.255002 | 799.657968 | 1071.88316 | 1353.3195 | 1636.90886 | 841.274512 | 1586.59041 |
| tRNA-Gly-TCC-2 | 998.163999 | 1432.9681 | 763.545425 | 848.82806 | 740.277426 | 1399.40302 | 1327.54199 | 1273.76269 | 1241.16285 | 1465.35593 |
| tRNA-Cys-GCA-14 | 956.789325 | 1259.04585 | 746.3871 | 870.513448 | 859.03851 | 967.672301 | 1271.6907 | 894.109879 | 1041.21868 | 1428.45848 |
| tRNA-Val-AAC-1 | 853.352642 | 1584.20484 | 879.364113 | 870.513448 | 597.764125 | 789.025107 | 897.916747 | 1094.94071 | 1078.94399 | 1112.19461 |
| tRNA-Ala-TGC-3 | 951.617491 | 714.593589 | 480.433076 | 771.380244 | 748.194832 | 1302.63579 | 1422.05954 | 938.127596 | 1301.52335 | 1101.65248 |
| tRNA-Val-AAC-4 | 703.36945 | 1251.48401 | 750.676681 | 978.94039 | 629.433747 | 893.235971 | 979.545542 | 1218.74054 | 1086.48906 | 1154.36312 |
| tRNA-Ser-GCT-4 | 630.963771 | 888.515838 | 806.441235 | 740.401118 | 633.39245 | 662.483345 | 661.622866 | 660.265757 | 494.201619 | 843.370319 |
| tRNA-Gly-GCC-5 | 548.214424 | 710.812671 | 514.749724 | 532.840972 | 787.78186 | 409.39982 | 747.547914 | 1199.48279 | 720.553506 | 848.641383 |
| tRNA-Ala-CGC-4 | 543.04259 | 722.155426 | 463.274752 | 731.10738 | 490.879149 | 364.738021 | 1061.17434 | 753.803406 | 694.145786 | 938.249479 |
| tRNA-Ser-CGA-1 | 641.30744 | 604.946954 | 673.464223 | 669.149127 | 621.516342 | 625.265179 | 618.660343 | 646.51022 | 724.326037 | 901.352028 |
| tRNA-Ser-CGA-4 | 713.713118 | 536.890421 | 600.541345 | 498.763933 | 676.938181 | 662.483345 | 777.621681 | 696.030152 | 735.643632 | 716.864771 |
| tRNA-Ser-GCT-1 | 527.527087 | 873.392164 | 810.730816 | 687.736603 | 562.135799 | 565.716115 | 644.437857 | 638.256898 | 497.974151 | 743.220093 |
| tRNA-Ser-GCT-2 | 475.808746 | 778.869203 | 759.255843 | 783.771894 | 645.268558 | 580.603381 | 597.179081 | 641.008006 | 441.386179 | 827.557125 |
| tRNA-Ala-TGC-6 | 605.1046 | 491.5194 | 411.79978 | 529.743059 | 597.764125 | 870.905071 | 923.694261 | 588.736967 | 762.051352 | 632.527739 |
| tRNA-Ser-CGA-2 | 605.1046 | 480.176645 | 660.59548 | 728.009467 | 621.516342 | 640.152446 | 588.586576 | 630.003576 | 652.64794 | 695.780513 |
| tRNA-Val-TAC-1 | 506.839751 | 608.727872 | 321.718578 | 439.903593 | 399.828984 | 729.476043 | 781.917933 | 687.77683 | 837.501981 | 853.912448 |
| tRNA-Ala-AGC-3 | 563.729927 | 710.812671 | 810.730816 | 570.015923 | 455.250823 | 617.821546 | 738.955409 | 610.745825 | 396.115802 | 579.817094 |
| tRNA-Trp-CCA-2 | 522.355253 | 514.204911 | 334.587321 | 424.41403 | 526.507474 | 632.708813 | 558.512809 | 596.990289 | 777.141477 | 706.322642 |
| tRNA-Gln-CTG-3 | 413.746735 | 608.727872 | 630.568412 | 675.344953 | 332.531036 | 565.716115 | 502.661528 | 522.710391 | 430.068585 | 564.003901 |
| tRNA-Ala-TGC-4 | 434.434072 | 366.749091 | 308.849835 | 381.043253 | 403.787687 | 684.814244 | 695.992885 | 423.670527 | 573.424779 | 627.256674 |
| tRNA-Asp-GTC-3 | 393.059399 | 680.565323 | 454.69559 | 381.043253 | 447.333418 | 454.061618 | 378.070209 | 437.426064 | 513.064276 | 411.14303 |
| tRNA-Phe-GAA-1 | 362.028393 | 597.385117 | 394.641455 | 387.239078 | 360.241956 | 424.287086 | 262.071395 | 635.505791 | 433.841116 | 490.208998 |
| tRNA-Arg-CCT-2 | 449.949575 | 291.130721 | 463.274752 | 412.022379 | 356.283253 | 409.39982 | 734.659157 | 434.674957 | 365.93555 | 347.890256 |
| tRNA-Arg-CCT-1 | 382.71573 | 306.254395 | 403.220617 | 504.959758 | 387.952876 | 424.287086 | 584.290324 | 349.39063 | 388.570739 | 437.498353 |
| tRNA-Cys-GCA-17 | 351.684725 | 517.985829 | 205.89989 | 346.966214 | 372.118064 | 431.730719 | 532.735295 | 332.883986 | 486.656556 | 574.54603 |
| tRNA-Gly-CCC-1 | 227.560704 | 654.098894 | 304.560254 | 244.735097 | 352.32455 | 297.745324 | 369.477705 | 437.426064 | 739.416163 | 458.582611 |
| tRNA-Thr-AGT-2 | 243.076207 | 412.120112 | 604.830926 | 285.007962 | 285.026602 | 491.279784 | 451.1065 | 338.3862 | 369.708082 | 558.732836 |
| tRNA-Glu-TTC-4 | 455.121409 | 472.614808 | 347.456064 | 353.162039 | 340.448442 | 409.39982 | 399.551471 | 332.883986 | 418.75099 | 421.685159 |
| tRNA-Leu-TAG-2 | 362.028393 | 385.653683 | 437.537266 | 319.085001 | 368.159362 | 409.39982 | 455.402752 | 426.421635 | 328.210236 | 426.956224 |
| tRNA-Pro-TGG-1 | 268.935378 | 389.434601 | 360.324807 | 278.812136 | 304.820117 | 513.610683 | 515.550285 | 354.892844 | 362.163019 | 495.480062 |
| tRNA-Ser-CGA-3 | 403.403067 | 264.664292 | 364.614388 | 402.728641 | 423.581201 | 394.512554 | 386.662714 | 332.883986 | 396.115802 | 442.769417 |
| tRNA-Gly-TCC-3 | 299.966383 | 483.957563 | 261.664443 | 346.966214 | 277.109197 | 357.294388 | 416.736481 | 357.643952 | 422.523522 | 484.937933 |

| | | | | | | | | | | |
|---|---|---|---|---|---|---|---|---|---|---|
| tRNA-Met-CAT-3 | 310.310051 | 241.978781 | 287.401929 | 303.595437 | 720.483912 | 372.181654 | 429.625238 | 343.888415 | 267.849733 | 332.077063 |
| tRNA-Leu-AAG-1 | 274.107212 | 468.833889 | 416.089361 | 288.105874 | 241.480872 | 320.076223 | 335.107686 | 478.692674 | 252.759607 | 405.871966 |
| tRNA-Trp-CCA-3 | 346.512891 | 313.816232 | 193.031147 | 340.770389 | 265.233088 | 468.948885 | 266.367648 | 385.155025 | 362.163019 | 469.12474 |
| tRNA-Leu-CAA-1 | 320.65372 | 298.692558 | 193.031147 | 223.049709 | 292.944008 | 521.054316 | 399.551471 | 203.581942 | 347.072893 | 400.600901 |
| tRNA-Met-CAT-6 | 279.279046 | 298.692558 | 223.058214 | 272.616311 | 213.769952 | 394.512554 | 416.736481 | 220.088586 | 441.386179 | 426.956224 |
| tRNA-Ser-GCT-5 | 191.357865 | 336.501743 | 454.69559 | 362.455777 | 245.439574 | 230.752626 | 352.292695 | 319.128449 | 169.763915 | 337.348127 |
| tRNA-Thr-AGT-5 | 129.295855 | 204.169597 | 364.614388 | 210.658059 | 364.200659 | 267.970791 | 416.736481 | 275.110732 | 339.52783 | 300.450676 |
| tRNA-Cys-GCA-5 | 263.763544 | 393.21552 | 210.189471 | 247.83301 | 197.935141 | 305.188957 | 274.960152 | 231.093015 | 279.167327 | 400.600901 |
| tRNA-Ala-CGC-2 | 237.904373 | 215.512352 | 214.479052 | 269.518399 | 237.522169 | 387.068921 | 369.477705 | 255.852981 | 298.029984 | 316.263869 |
| tRNA-Thr-TGT-2 | 181.014197 | 279.787966 | 295.981092 | 260.224661 | 229.604763 | 260.527158 | 283.552657 | 209.084156 | 237.669481 | 358.432385 |
| tRNA-Pro-CGG-2 | 212.045202 | 347.844498 | 223.058214 | 201.364321 | 273.150494 | 387.068921 | 257.775143 | 184.32419 | 199.944167 | 284.637483 |
| tRNA-Gln-CTG-6 | 186.186031 | 162.579494 | 120.108269 | 226.147622 | 201.893843 | 193.53446 | 257.775143 | 239.346337 | 256.532138 | 237.197902 |
| tRNA-Gln-CTG-5 | 237.904373 | 196.60776 | 154.424917 | 232.343447 | 170.224221 | 133.985396 | 227.701376 | 198.079727 | 192.399104 | 284.637483 |
| tRNA-Ile-AAT-7 | 165.498694 | 139.893983 | 201.610309 | 111.524855 | 265.233088 | 267.970791 | 283.552657 | 151.310903 | 207.489229 | 226.655773 |
| tRNA-Glu-TTC-3 | 160.32686 | 234.416945 | 167.29366 | 188.97267 | 154.38941 | 253.083525 | 171.850095 | 165.066439 | 173.536446 | 142.318741 |
| tRNA-Asn-GTT-2 | 103.436684 | 226.855108 | 171.583241 | 182.776845 | 142.513301 | 171.203561 | 176.146348 | 165.066439 | 188.626572 | 163.402999 |
| tRNA-Gln-CTG-4 | 129.295855 | 173.922249 | 141.556174 | 148.699806 | 150.430707 | 148.872662 | 206.220114 | 217.337478 | 128.266069 | 242.468967 |
| tRNA-Lys-TTT-5 | 201.701533 | 151.236738 | 81.5020397 | 148.699806 | 225.64606 | 200.978093 | 120.295067 | 148.559795 | 196.171635 | 179.216193 |
| tRNA-Thr-TGT-4 | 155.155026 | 94.5229615 | 94.3707828 | 108.426942 | 162.306815 | 186.090827 | 163.25759 | 82.5332196 | 162.218852 | 289.908547 |
| tRNA-Ala-AGC-1 | 118.952186 | 139.893983 | 240.216538 | 58.8603399 | 51.4631366 | 111.654496 | 227.701376 | 162.315332 | 169.763915 | 200.300451 |
| tRNA-Thr-TGT-5 | 124.124021 | 136.113065 | 111.529107 | 86.7415535 | 142.513301 | 208.421726 | 206.220114 | 143.057581 | 132.038601 | 189.758322 |
| tRNA-Thr-TGT-3 | 139.639523 | 113.427554 | 90.0812018 | 130.11233 | 102.926273 | 171.203561 | 180.4426 | 104.542078 | 184.854041 | 152.86087 |
| tRNA-Leu-CAA-4 | 175.842362 | 113.427554 | 55.7645535 | 68.1540778 | 142.513301 | 178.647194 | 171.850095 | 170.568654 | 128.266069 | 158.131935 |
| tRNA-Leu-TAA-2 | 118.952186 | 117.208472 | 150.135336 | 105.329029 | 182.100329 | 119.098129 | 115.998814 | 143.057581 | 169.763915 | 126.505548 |
| tRNA-Thr-AGT-6 | 134.467689 | 109.646635 | 167.29366 | 108.426942 | 98.9675703 | 133.985396 | 146.072581 | 143.057581 | 139.583663 | 137.047677 |
| tRNA-Ile-AAT-5 | 113.780352 | 60.4946954 | 154.424917 | 89.8394662 | 213.769952 | 111.654496 | 124.591319 | 85.2843269 | 124.493538 | 158.131935 |
| tRNA-Leu-CAA-2 | 149.983192 | 79.3992877 | 55.7645535 | 74.349903 | 98.9675703 | 178.647194 | 171.850095 | 107.293186 | 154.673789 | 94.8791608 |
| tRNA-Val-TAC-2 | 118.952186 | 83.1802061 | 60.0541345 | 99.133204 | 83.1327591 | 178.647194 | 150.368833 | 96.2887562 | 158.446321 | 121.234483 |
| tRNA-Ala-AGC-5 | 651.651108 | 26.4664292 | 42.8958104 | 37.1749515 | 35.6283253 | 74.4363309 | 55.8512809 | 38.5155025 | 52.8154402 | 52.7106449 |
| tRNA-Arg-CCT-3 | 77.5775129 | 113.427554 | 132.977012 | 86.7415535 | 75.2153534 | 104.210863 | 124.591319 | 104.542078 | 120.721006 | 79.0659674 |
| tRNA-Ile-AAT-12 | 82.7493471 | 56.7137769 | 72.9228776 | 89.8394662 | 55.4218394 | 200.978093 | 94.5175523 | 82.5332196 | 105.63088 | 147.589806 |
| tRNA-Cys-GCA-19 | 139.639523 | 158.798575 | 38.6062293 | 52.6645147 | 59.3805422 | 59.5490647 | 120.295067 | 85.2843269 | 94.3132861 | 152.86087 |
| tRNA-Asn-GTT-4 | 67.2338445 | 90.7420431 | 107.239526 | 65.0561652 | 126.67849 | 126.541763 | 111.702562 | 101.790971 | 86.7682232 | 68.5238384 |
| tRNA-Glu-TTC-14 | 103.436684 | 136.113065 | 124.39785 | 111.524855 | 134.595896 | 126.541763 | 51.5550285 | 44.0177171 | 64.1330345 | 36.8974514 |
| tRNA-Ile-AAT-8 | 36.2028393 | 34.0282661 | 120.108269 | 92.9373788 | 150.430707 | 81.879964 | 107.406309 | 46.7688245 | 128.266069 | 84.3370319 |
| tRNA-Tyr-GTA-2 | 62.0620103 | 90.7420431 | 51.4749724 | 27.8812136 | 63.339245 | 126.541763 | 197.627609 | 27.5110732 | 60.3605031 | 131.776612 |
| tRNA-Val-AAC-3 | 46.5465077 | 132.332146 | 85.7916207 | 108.426942 | 63.339245 | 37.2181654 | 77.3325428 | 77.031005 | 86.7682232 | 94.8791608 |
| tRNA-Arg-TCG-5 | 56.8901761 | 102.084798 | 90.0812018 | 74.349903 | 67.2979478 | 81.879964 | 68.7400381 | 101.790971 | 49.0429088 | 115.963419 |
| tRNA-Ile-AAT-1 | 72.4056787 | 41.5901031 | 94.3707828 | 74.349903 | 106.884976 | 44.6617985 | 128.887571 | 66.0265757 | 56.5879717 | 110.692354 |
| tRNA-Ile-AAT-6 | 98.2648496 | 49.15194 | 8.57916207 | 43.3707768 | 59.3805422 | 111.654496 | 115.998814 | 82.5332196 | 90.5407547 | 105.42129 |
| tRNA-Arg-TCG-4 | 46.5465077 | 75.6183692 | 72.9228776 | 74.349903 | 75.2153534 | 59.5490647 | 94.5175523 | 74.2798977 | 90.5407547 | 84.3370319 |
| tRNA-Trp-CCA-1 | 98.2648496 | 75.6183692 | 34.3166483 | 80.5457283 | 67.2979478 | 52.1054316 | 68.7400381 | 107.293186 | 75.4506289 | 84.3370319 |

| | | | | | | | | | | |
|---|---|---|---|---|---|---|---|---|---|---|
| tRNA-Asn-GTT-6 | 41.3746735 | 86.9611246 | 42.8958104 | 92.9373788 | 102.926273 | 66.9926978 | 77.3325428 | 60.5243611 | 82.9956918 | 63.2527739 |
| tRNA-Arg-TCT-3 | 77.5775129 | 37.8091846 | 64.3437155 | 74.349903 | 75.2153534 | 89.3235971 | 77.3325428 | 66.0265757 | 52.8154402 | 94.8791608 |
| tRNA-Lys-CTT-11 | 67.2338445 | 86.9611246 | 72.9228776 | 74.349903 | 27.7109197 | 37.2181654 | 73.0362904 | 49.5199318 | 109.403412 | 84.3370319 |
| tRNA-Arg-TCG-2 | 41.3746735 | 60.4946954 | 60.0541345 | 49.566602 | 67.2979478 | 66.9926978 | 77.3325428 | 104.542078 | 64.1330345 | 52.7106449 |
| tRNA-Thr-AGT-3 | 36.2028393 | 41.5901031 | 60.0541345 | 68.1540778 | 47.5044337 | 66.9926978 | 55.8512809 | 63.2754684 | 79.2231603 | 73.7949029 |
| tRNA-Phe-GAA-2 | 20.6873368 | 68.0565323 | 42.8958104 | 61.9582525 | 35.6283253 | 52.1054316 | 51.5550285 | 71.5287903 | 45.2703773 | 110.692354 |
| tRNA-Gln-TTG-2 | 62.0620103 | 18.9045923 | 30.0270673 | 58.8603399 | 51.4631366 | 59.5490647 | 64.4437857 | 49.5199318 | 64.1330345 | 89.6080964 |
| tRNA-Asn-GTT-1 | 20.6873368 | 71.8374508 | 72.9228776 | 46.4686894 | 23.7522169 | 44.6617985 | 68.7400381 | 74.2798977 | 56.5879717 | 52.7106449 |
| tRNA-Ala-TGC-2 | 51.7183419 | 52.9328584 | 25.7374862 | 49.566602 | 39.5870281 | 52.1054316 | 85.9250476 | 33.0132878 | 86.7682232 | 47.4395804 |
| tRNA-Tyr-GTA-5-1 | 25.859171 | 41.5901031 | 30.0270673 | 12.3916505 | 55.4218394 | 74.4363309 | 90.2213 | 49.5199318 | 52.8154402 | 89.6080964 |
| tRNA-Ile-TAT-2-1 | 67.2338445 | 52.9328584 | 21.4479052 | 15.4895631 | 11.8761084 | 37.2181654 | 68.7400381 | 44.0177171 | 75.4506289 | 94.8791608 |
| tRNA-Arg-TCT-5 | 15.5155026 | 37.8091846 | 42.8958104 | 40.2728641 | 59.3805422 | 52.1054316 | 77.3325428 | 49.5199318 | 37.7253144 | 26.3553225 |
| tRNA-Leu-AAG-4 | 51.7183419 | 68.0565323 | 34.3166483 | 34.0770389 | 39.5870281 | 22.3308993 | 38.6662714 | 46.7688245 | 41.4978459 | 52.7106449 |
| tRNA-Cys-GCA-11 | 41.3746735 | 64.2756138 | 34.3166483 | 40.2728641 | 63.339245 | 37.2181654 | 34.370019 | 22.0088586 | 37.7253144 | 42.1685159 |
| tRNA-Cys-GCA-15 | 31.0310051 | 60.4946954 | 4.28958104 | 24.783301 | 19.7935141 | 29.7745324 | 34.370019 | 41.2666098 | 60.3605031 | 110.692354 |
| tRNA-Ile-GAT-1 | 15.5155026 | 18.9045923 | 25.7374862 | 27.8812136 | 63.339245 | 66.9926978 | 81.6287952 | 27.5110732 | 26.4077201 | 52.7106449 |
| tRNA-Val-TAC-3 | 36.2028393 | 52.9328584 | 17.1583241 | 27.8812136 | 19.7935141 | 44.6617985 | 42.9625238 | 49.5199318 | 33.952783 | 57.9817094 |
| tRNA-Ser-GCT-6 | 36.2028393 | 68.0565323 | 47.1853914 | 18.5874758 | 31.6696225 | 29.7745324 | 17.1850095 | 49.5199318 | 45.2703773 | 36.8974514 |
| tRNA-Thr-CGT-3 | 41.3746735 | 18.9045923 | 25.7374862 | 40.2728641 | 43.5457309 | 52.1054316 | 38.6662714 | 41.2666098 | 41.4978459 | 36.8974514 |
| tRNA-Thr-TGT-6 | 10.3436684 | 22.6855108 | 34.3166483 | 34.0770389 | 23.7522169 | 44.6617985 | 68.7400381 | 33.0132878 | 52.8154402 | 31.6263869 |
| tRNA-Tyr-GTA-5-5 | 31.0310051 | 34.0282661 | 21.4479052 | 3.09791263 | 27.7109197 | 66.9926978 | 51.5550285 | 22.0088586 | 37.7253144 | 52.7106449 |
| tRNA-Ile-TAT-3 | 31.0310051 | 18.9045923 | 8.57916207 | 15.4895631 | 27.7109197 | 29.7745324 | 55.8512809 | 41.2666098 | 33.952783 | 79.0659674 |
| tRNA-Cys-GCA-7 | 41.3746735 | 49.15194 | 25.7374862 | 40.2728641 | 31.6696225 | 29.7745324 | 34.370019 | 24.7599659 | 15.0901258 | 47.4395804 |
| tRNA-Ile-AAT-4 | 31.0310051 | 15.1236738 | 21.4479052 | 24.783301 | 35.6283253 | 44.6617985 | 60.1475333 | 16.5066439 | 30.1802516 | 21.084258 |
| tRNA-Ile-AAT-2 | 25.859171 | 11.3427554 | 21.4479052 | 3.09791263 | 27.7109197 | 52.1054316 | 47.2587762 | 35.7643952 | 26.4077201 | 42.1685159 |
| tRNA-Ile-TAT-1 | 36.2028393 | 41.5901031 | 21.4479052 | 12.3916505 | 15.8348112 | 44.6617985 | 17.1850095 | 11.0044293 | 18.8626572 | 73.7949029 |
| tRNA-Arg-TCT-2 | 15.5155026 | 26.4664292 | 12.8687431 | 55.7624273 | 31.6696225 | 7.44363309 | 34.370019 | 30.2621805 | 33.952783 | 15.8131935 |
| tRNA-Leu-CAA-6 | 25.859171 | 26.4664292 | 21.4479052 | 24.783301 | 31.6696225 | 22.3308993 | 38.6662714 | 16.5066439 | 33.952783 | 21.084258 |
| tRNA-Ala-AGC-9 | 15.5155026 | 18.9045923 | 25.7374862 | 18.5874758 | 31.6696225 | 7.44363309 | 25.7775143 | 38.5155025 | 18.8626572 | 15.8131935 |
| tRNA-Tyr-GTA-5-4 | 41.3746735 | 7.56183692 | 17.1583241 | 15.4895631 | 7.91740562 | 29.7745324 | 21.4812619 | 8.25332196 | 30.1802516 | 36.8974514 |
| tRNA-Cys-GCA-8 | 15.5155026 | 18.9045923 | 4.28958104 | 9.29373788 | 19.7935141 | 14.8872662 | 25.7775143 | 27.5110732 | 18.8626572 | 31.6263869 |
| tRNA-Val-TAC-4 | 15.5155026 | 26.4664292 | 4.28958104 | 15.4895631 | 19.7935141 | 7.44363309 | 17.1850095 | 35.7643952 | 11.3175943 | 26.3553225 |
| tRNA-Cys-GCA-3 | 10.3436684 | 34.0282661 | 17.1583241 | 12.3916505 | 11.8761084 | 14.8872662 | 12.8887571 | 16.5066439 | 11.3175943 | 36.8974514 |
| tRNA-Tyr-GTA-4 | 15.5155026 | 18.9045923 | 4.28958104 | 21.6853884 | 23.7522169 | 14.8872662 | 21.4812619 | 11.0044293 | 30.1802516 | 15.8131935 |
| tRNA-Ala-TGC-1 | 0 | 18.9045923 | 12.8687431 | 18.5874758 | 7.91740562 | 22.3308993 | 17.1850095 | 13.7555366 | 26.4077201 | 36.8974514 |
| tRNA-Ala-AGC-15 | 5.17183419 | 11.3427554 | 21.4479052 | 6.19582525 | 11.8761084 | 22.3308993 | 8.59250476 | 22.0088586 | 22.6351887 | 42.1685159 |
| tRNA-Trp-CCA-5 | 10.3436684 | 34.0282661 | 17.1583241 | 3.09791263 | 19.7935141 | 7.44363309 | 21.4812619 | 13.7555366 | 7.54506289 | 36.8974514 |
| tRNA-Cys-GCA-23 | 5.17183419 | 22.6855108 | 0 | 18.5874758 | 15.8348112 | 22.3308993 | 4.29625238 | 13.7555366 | 18.8626572 | 31.6263869 |
| tRNA-Tyr-GTA-1 | 20.6873368 | 7.56183692 | 12.8687431 | 9.29373788 | 27.7109197 | 29.7745324 | 8.59250476 | 5.50221464 | 22.6351887 | 5.27106449 |
| tRNA-Met-CAT-2 | 0 | 3.78091846 | 0 | 58.8603399 | 11.8761084 | 0 | 0 | 8.25332196 | 3.77253144 | 57.9817094 |
| tRNA-Asn-GTT-24 | 15.5155026 | 15.1236738 | 0 | 9.29373788 | 23.7522169 | 14.8872662 | 12.8887571 | 11.0044293 | 22.6351887 | 15.8131935 |
| tRNA-Ile-AAT-3 | 10.3436684 | 7.56183692 | 4.28958104 | 6.19582525 | 27.7109197 | 14.8872662 | 12.8887571 | 11.0044293 | 26.4077201 | 15.8131935 |

| | | | | | | | | | | |
|---|---|---|---|---|---|---|---|---|---|---|
| tRNA-Asn-GTT-5 | 15.5155026 | 26.4664292 | 21.4479052 | 6.19582525 | 7.91740562 | 7.44363309 | 8.59250476 | 13.7555366 | 15.0901258 | 10.542129 |
| tRNA-Thr-CGT-1 | 10.3436684 | 22.6855108 | 12.8687431 | 15.4895631 | 15.8348112 | 7.44363309 | 8.59250476 | 5.50221464 | 7.54506289 | 21.084258 |
| tRNA-Gln-TTG-4 | 15.5155026 | 11.3427554 | 8.57916207 | 0 | 15.8348112 | 22.3308993 | 25.7775143 | 0 | 7.54506289 | 10.542129 |
| tRNA-Thr-TGT-1 | 0 | 18.9045923 | 12.8687431 | 12.3916505 | 3.95870281 | 0 | 12.8887571 | 11.0044293 | 11.3175943 | 31.6263869 |
| tRNA-Gln-CTG-7 | 0 | 11.3427554 | 21.4479052 | 18.5874758 | 0 | 0 | 12.8887571 | 13.7555366 | 15.0901258 | 15.8131935 |
| tRNA-Phe-GAA-3 | 0 | 11.3427554 | 4.28958104 | 9.29373788 | 7.91740562 | 14.8872662 | 17.1850095 | 8.25332196 | 3.77253144 | 21.084258 |
| tRNA-Leu-TAA-4 | 0 | 18.9045923 | 8.57916207 | 15.4895631 | 11.8761084 | 7.44363309 | 12.8887571 | 11.0044293 | 3.77253144 | 5.27106449 |
| tRNA-Gly-TCC-4 | 0 | 11.3427554 | 0 | 6.19582525 | 11.8761084 | 7.44363309 | 4.29625238 | 13.7555366 | 30.1802516 | 5.27106449 |
| tRNA-Asn-GTT-7 | 0 | 18.9045923 | 0 | 9.29373788 | 3.95870281 | 14.8872662 | 4.29625238 | 0 | 7.54506289 | 26.3553225 |
| tRNA-Leu-TAA-3 | 5.17183419 | 11.3427554 | 17.1583241 | 6.19582525 | 3.95870281 | 14.8872662 | 4.29625238 | 8.25332196 | 7.54506289 | 5.27106449 |
| tRNA-Tyr-GTA-8 | 10.3436684 | 7.56183692 | 4.28958104 | 9.29373788 | 11.8761084 | 0 | 8.59250476 | 2.75110732 | 22.6351887 | 5.27106449 |
| tRNA-Tyr-GTA-9 | 5.17183419 | 7.56183692 | 0 | 9.29373788 | 3.95870281 | 14.8872662 | 8.59250476 | 0 | 15.0901258 | 5.27106449 |
| tRNA-Cys-GCA-13 | 5.17183419 | 15.1236738 | 8.57916207 | 6.19582525 | 3.95870281 | 0 | 4.29625238 | 2.75110732 | 22.6351887 | 0 |
| tRNA-Ala-AGC-6 | 15.5155026 | 3.78091846 | 4.28958104 | 3.09791263 | 11.8761084 | 7.44363309 | 12.8887571 | 0 | 3.77253144 | 5.27106449 |
| tRNA-Asn-GTT-26 | 0 | 7.56183692 | 8.57916207 | 3.09791263 | 3.95870281 | 7.44363309 | 12.8887571 | 8.25332196 | 3.77253144 | 10.542129 |
| tRNA-Asn-GTT-9 | 0 | 15.1236738 | 4.28958104 | 12.3916505 | 3.95870281 | 14.8872662 | 0 | 0 | 3.77253144 | 10.542129 |
| tRNA-Lys-CTT-8 | 10.3436684 | 3.78091846 | 0 | 6.19582525 | 3.95870281 | 22.3308993 | 4.29625238 | 5.50221464 | 0 | 5.27106449 |
| tRNA-Cys-GCA-6 | 5.17183419 | 7.56183692 | 0 | 3.09791263 | 0 | 0 | 12.8887571 | 5.50221464 | 3.77253144 | 21.084258 |
| tRNA-Lys-TTT-6 | 5.17183419 | 3.78091846 | 0 | 3.09791263 | 7.91740562 | 0 | 12.8887571 | 8.25332196 | 11.3175943 | 5.27106449 |
| tRNA-Leu-CAA-5 | 5.17183419 | 3.78091846 | 4.28958104 | 3.09791263 | 7.91740562 | 0 | 8.59250476 | 8.25332196 | 0 | 15.8131935 |
| tRNA-Tyr-GTA-3 | 5.17183419 | 3.78091846 | 0 | 3.09791263 | 7.91740562 | 14.8872662 | 4.29625238 | 2.75110732 | 3.77253144 | 10.542129 |
| tRNA-Lys-CTT-7 | 10.3436684 | 3.78091846 | 0 | 9.29373788 | 0 | 7.44363309 | 8.59250476 | 2.75110732 | 7.54506289 | 5.27106449 |
| tRNA-Cys-GCA-18 | 0 | 7.56183692 | 0 | 0 | 11.8761084 | 7.44363309 | 4.29625238 | 11.0044293 | 7.54506289 | 5.27106449 |
| tRNA-Cys-GCA-21 | 5.17183419 | 11.3427554 | 4.28958104 | 6.19582525 | 0 | 0 | 4.29625238 | 5.50221464 | 0 | 15.8131935 |
| tRNA-Gly-GCC-4 | 5.17183419 | 0 | 4.28958104 | 6.19582525 | 7.91740562 | 7.44363309 | 4.29625238 | 2.75110732 | 7.54506289 | 5.27106449 |
| tRNA-Ala-TGC-5 | 5.17183419 | 3.78091846 | 0 | 0 | 0 | 14.8872662 | 0 | 8.25332196 | 7.54506289 | 5.27106449 |
| tRNA-Tyr-GTA-5-3 | 0 | 0 | 0 | 0 | 3.95870281 | 7.44363309 | 12.8887571 | 5.50221464 | 7.54506289 | 0 |
| tRNA-Cys-GCA-12 | 5.17183419 | 7.56183692 | 8.57916207 | 3.09791263 | 3.95870281 | 0 | 0 | 0 | 7.54506289 | 0 |
| tRNA-Thr-AGT-4 | 5.17183419 | 0 | 0 | 3.09791263 | 0 | 0 | 4.29625238 | 8.25332196 | 7.54506289 | 5.27106449 |
| tRNA-Cys-GCA-10 | 0 | 0 | 4.28958104 | 0 | 0 | 7.44363309 | 4.29625238 | 2.75110732 | 3.77253144 | 10.542129 |
| tRNA-Tyr-GTA-7 | 0 | 0 | 4.28958104 | 0 | 3.95870281 | 0 | 4.29625238 | 2.75110732 | 11.3175943 | 5.27106449 |
| tRNA-Lys-TTT-4 | 0 | 7.56183692 | 4.28958104 | 0 | 3.95870281 | 0 | 0 | 2.75110732 | 7.54506289 | 5.27106449 |
| tRNA-Asn-GTT-11 | 0 | 3.78091846 | 8.57916207 | 6.19582525 | 3.95870281 | 0 | 0 | 2.75110732 | 3.77253144 | 0 |
| tRNA-Gly-CCC-3 | 0 | 0 | 8.57916207 | 6.19582525 | 0 | 0 | 0 | 2.75110732 | 3.77253144 | 5.27106449 |
| tRNA-Cys-GCA-9 | 0 | 3.78091846 | 0 | 3.09791263 | 3.95870281 | 0 | 4.29625238 | 0 | 0 | 10.542129 |
| tRNA-Asn-GTT-8 | 10.3436684 | 0 | 0 | 3.09791263 | 0 | 0 | 0 | 0 | 0 | 5.27106449 |
| tRNA-Tyr-ATA-1 | 0 | 3.78091846 | 4.28958104 | 3.09791263 | 3.95870281 | 0 | 0 | 2.75110732 | 0 | 0 |
| tRNA-Lys-TTT-11 | 0 | 3.78091846 | 0 | 0 | 0 | 0 | 8.59250476 | 0 | 0 | 5.27106449 |
| tRNA-Lys-CTT-10 | 5.17183419 | 0 | 4.28958104 | 0 | 0 | 0 | 4.29625238 | 0 | 3.77253144 | 0 |
| tRNA-Met-CAT-4 | 0 | 0 | 0 | 6.19582525 | 0 | 7.44363309 | 0 | 0 | 3.77253144 | 0 |
| tRNA-Lys-TTT-1 | 0 | 3.78091846 | 0 | 0 | 0 | 7.44363309 | 4.29625238 | 0 | 0 | 0 |
| tRNA-Phe-GAA-4 | 5.17183419 | 3.78091846 | 0 | 0 | 0 | 0 | 0 | 2.75110732 | 3.77253144 | 0 |

| | | | | | | | | | | |
|---|---|---|---|---|---|---|---|---|---|---|
| tRNA-Ile-TAT-2 | 0 | 0 | 0 | 0 | 0 | 0 | 8.59250476 | 2.75110732 | 3.77253144 | 0 |
| tRNA-Lys-CTT-9 | 0 | 0 | 0 | 0 | 0 | 7.44363309 | 0 | 5.50221464 | 0 | 0 |
| tRNA-Lys-TTT-2 | 5.17183419 | 3.78091846 | 0 | 3.09791263 | 0 | 0 | 0 | 0 | 0 | 0 |
| tRNA-Cys-GCA-20 | 0 | 0 | 4.28958104 | 0 | 0 | 0 | 4.29625238 | 2.75110732 | 0 | 0 |
| tRNA-Ile-TAT-2-2 | 0 | 0 | 0 | 0 | 0 | 7.44363309 | 0 | 0 | 3.77253144 | 0 |
| tRNA-Tyr-GTA-5-2 | 0 | 0 | 0 | 0 | 7.91740562 | 0 | 0 | 2.75110732 | 0 | 0 |
| tRNA-Arg-CCT-5 | 0 | 0 | 0 | 3.09791263 | 3.95870281 | 0 | 0 | 2.75110732 | 0 | 0 |
| tRNA-Arg-TCG-6 | 0 | 0 | 0 | 0 | 0 | 7.44363309 | 0 | 0 | 0 | 0 |
| tRNA-Cys-GCA-16 | 0 | 0 | 0 | 0 | 0 | 7.44363309 | 0 | 0 | 0 | 0 |
| tRNA-Asn-GTT-10 | 0 | 0 | 0 | 0 | 0 | 0 | 0 | 0 | 0 | 5.27106449 |
| tRNA-Tyr-GTA-6 | 0 | 0 | 0 | 0 | 0 | 0 | 4.29625238 | 0 | 0 | 0 |
| tRNA-Ala-TGC-7 | 0 | 0 | 0 | 0 | 3.95870281 | 0 | 0 | 0 | 0 | 0 |
| tRNA-Asn-GTT-25 | 0 | 0 | 0 | 0 | 3.95870281 | 0 | 0 | 0 | 0 | 0 |
| tRNA-Thr-CGT-5 | 0 | 3.78091846 | 0 | 0 | 0 | 0 | 0 | 0 | 0 | 0 |
| tRNA-Phe-GAA-6 | 0 | 0 | 0 | 3.09791263 | 0 | 0 | 0 | 0 | 0 | 0 |
| tRNA-Lys-TTT-7 | 0 | 0 | 0 | 3.09791263 | 0 | 0 | 0 | 0 | 0 | 0 |
| tRNA-Asn-GTT-12 | 0 | 0 | 0 | 3.09791263 | 0 | 0 | 0 | 0 | 0 | 0 |
| tRNA-Val-AAC-6 | 0 | 0 | 0 | 3.09791263 | 0 | 0 | 0 | 0 | 0 | 0 |
| tRNA-Cys-GCA-22 | 0 | 0 | 0 | 0 | 0 | 0 | 0 | 2.75110732 | 0 | 0 |
| tRNA-Met-CAT-7 | 0 | 0 | 0 | 0 | 0 | 0 | 0 | 0 | 0 | 0 |
| tRNA-Met-CAT-5 | 0 | 0 | 0 | 0 | 0 | 0 | 0 | 0 | 0 | 0 |
| tRNA-Ala-AGC-24 | 0 | 0 | 0 | 0 | 0 | 0 | 0 | 0 | 0 | 0 |
| tRNA-Ala-AGC-16 | 0 | 0 | 0 | 0 | 0 | 0 | 0 | 0 | 0 | 0 |
| tRNA-Asn-GTT-27 | 0 | 0 | 0 | 0 | 0 | 0 | 0 | 0 | 0 | 0 |
| tRNA-Ala-AGC-14 | 0 | 0 | 0 | 0 | 0 | 0 | 0 | 0 | 0 | 0 |
| tRNA-Ala-AGC-13 | 0 | 0 | 0 | 0 | 0 | 0 | 0 | 0 | 0 | 0 |
| tRNA-Ala-AGC-12 | 0 | 0 | 0 | 0 | 0 | 0 | 0 | 0 | 0 | 0 |
| tRNA-Ala-AGC-10 | 0 | 0 | 0 | 0 | 0 | 0 | 0 | 0 | 0 | 0 |
| tRNA-Ile-AAT-9 | 0 | 0 | 0 | 0 | 0 | 0 | 0 | 0 | 0 | 0 |

#### Table S3. tRNA isodecoder expression [tpm].

| isodecoder | HD1 | HD2 | HD3 | HD4 | HD5 | HD6 | HD7 | HD8 | HD9 | HD10 |
|---|---|---|---|---|---|---|---|---|---|---|
| tRNA-Ser-AGA | 142085.801 | 107192.819 | 125435.929 | 101100.379 | 139674.911 | 88035.8485 | 126146.562 | 143387.714 | 123444.774 | 85217.2996 |
| tRNA-Ser-TGA | 116578.314 | 90889.4989 | 107273.843 | 89374.7793 | 121995.345 | 77257.4678 | 115668.003 | 127362.513 | 111153.866 | 73858.1557 |
| tRNA-Asp-GTC | 80871.9712 | 60574.0947 | 110924.276 | 108786.3 | 105535.058 | 89517.1315 | 59357.0229 | 60562.8766 | 69142.9563 | 56300.2398 |
| tRNA-Gly-GCC | 63882.4959 | 116766.105 | 64867.0444 | 72828.8279 | 62036.8318 | 53117.7657 | 78346.4584 | 94288.7012 | 71591.3292 | 95132.1719 |
| tRNA-Glu-CTC | 97514.9337 | 61118.5469 | 90788.9826 | 84219.8527 | 85135.8627 | 111974.573 | 51988.95 | 36650.2517 | 64823.4078 | 58071.3175 |
| tRNA-Ala-AGC | 64441.054 | 78631.7612 | 70168.9666 | 67438.46 | 47607.36 | 65295.5495 | 75931.9645 | 67245.3162 | 66223.017 | 78465.066 |
| tRNA-Arg-TCT | 52307.931 | 57742.1867 | 45606.8256 | 59411.7684 | 71937.5475 | 46768.3467 | 59924.1282 | 60428.0723 | 60692.4859 | 62024.6159 |
| tRNA-Leu-CAG | 36637.2734 | 53329.8549 | 59625.1764 | 46927.1805 | 40600.456 | 38319.8231 | 47980.5466 | 57013.9481 | 39075.8807 | 59399.6258 |
| tRNA-Lys-CTT | 54624.9127 | 36950.9161 | 41922.0755 | 52218.4152 | 40473.7776 | 57636.051 | 42945.3388 | 33753.3357 | 54784.7016 | 48694.0938 |
| tRNA-Arg-ACG | 31770.5774 | 34739.0788 | 25475.8218 | 31731.919 | 33383.7408 | 42562.694 | 38219.4612 | 37258.2464 | 41765.6956 | 44967.4512 |

| | | | | | | | | | | |
|---|---|---|---|---|---|---|---|---|---|---|
| tRNA-iMet-CAT | 26515.9939 | 35045.3332 | 24647.9326 | 31849.6397 | 24785.4383 | 45279.6201 | 37179.7681 | 42994.3052 | 42720.1461 | 49316.0794 |
| tRNA-Gln-CTG | 20273.59 | 21369.7511 | 19985.158 | 24851.4551 | 18558.3988 | 21050.5944 | 28015.8618 | 26528.9279 | 23242.5662 | 27984.0814 |
| tRNA-Pro-AGG | 21204.5202 | 21725.1575 | 20512.7765 | 18695.9027 | 18867.1776 | 28181.5949 | 19895.9448 | 13780.2966 | 17410.2326 | 24515.7209 |
| tRNA-Val-CAC | 14750.0711 | 25785.8639 | 18488.0943 | 19826.6408 | 12806.4036 | 16100.5784 | 19552.2446 | 24107.9535 | 21371.3906 | 23329.7314 |
| tRNA-Ser-GCT | 14812.1331 | 23097.6309 | 22395.9026 | 20845.8541 | 17786.4517 | 18385.7737 | 20527.4939 | 17563.0691 | 15739.0012 | 22365.1266 |
| tRNA-Gly-CCC | 19022.0062 | 21120.2105 | 14267.1465 | 18940.6378 | 14421.5543 | 20179.6893 | 17249.4533 | 17029.3543 | 20473.5281 | 18870.4109 |
| tRNA-SeC-TCA | 13932.9213 | 15724.8399 | 14790.4754 | 19185.3729 | 19239.2957 | 16160.1274 | 15161.4746 | 16223.2799 | 17900.6617 | 13525.5515 |
| tRNA-Leu-AAG | 13689.8451 | 21067.2777 | 14683.2359 | 15582.5005 | 13629.8138 | 15170.1242 | 16265.6115 | 16060.9645 | 14286.5766 | 18522.5206 |
| tRNA-Pro-CGG | 15272.4264 | 15736.1826 | 14249.9882 | 14163.6565 | 14061.3124 | 20648.6382 | 14740.4419 | 10467.9634 | 12426.7186 | 18617.3998 |
| tRNA-Glu-TTC | 11212.5365 | 7554.27508 | 15185.1169 | 10213.8179 | 10819.1348 | 16457.8728 | 8961.98246 | 8853.06336 | 9699.17834 | 9888.51699 |
| tRNA-Arg-CCT | 11709.0326 | 8348.26796 | 11345.9418 | 12050.8801 | 11745.4712 | 14075.9102 | 9808.34418 | 7249.16779 | 10423.5044 | 10378.726 |
| tRNA-Pro-TGG | 10948.773 | 10541.2007 | 8094.43942 | 9371.1857 | 10391.5949 | 13309.216 | 10783.5935 | 8445.89947 | 9816.12682 | 11759.7449 |
| tRNA-Ala-CGC | 5968.29666 | 6480.49424 | 6078.33633 | 8209.46846 | 5526.34913 | 7570.17485 | 10469.967 | 8330.35297 | 7579.01567 | 8328.2819 |
| tRNA-Arg-CCG | 6263.09121 | 6064.59321 | 5923.91141 | 6855.68064 | 6979.19306 | 8597.39622 | 8227.32331 | 6533.87989 | 6922.5952 | 7990.93377 |
| tRNA-His-GTG | 4685.68178 | 8253.745 | 7142.15243 | 8274.52463 | 7082.11933 | 5835.80834 | 5847.19949 | 6522.87546 | 6526.4794 | 6309.4642 |
| tRNA-Cys-GCA | 5120.11585 | 9879.53994 | 3543.19394 | 4594.20443 | 4675.22802 | 5158.43773 | 6715.04247 | 4993.25979 | 6730.1961 | 8918.64112 |
| tRNA-Gln-TTG | 6076.90517 | 4015.33541 | 2612.35485 | 4442.40671 | 4386.24272 | 7942.35651 | 8390.5809 | 5983.65842 | 7333.80113 | 7400.57455 |
| tRNA-Val-AAC | 4556.38592 | 7501.34223 | 4225.23732 | 5666.08219 | 3214.46668 | 4853.24877 | 6070.60461 | 6902.52827 | 7330.0286 | 6836.57065 |
| tRNA-Arg-TCG | 3351.34856 | 3051.2012 | 2642.38192 | 3801.13879 | 3436.15404 | 4711.81974 | 5469.12928 | 3829.54139 | 4843.93037 | 5228.89598 |
| tRNA-Leu-TAG | 2704.86928 | 4215.72408 | 3526.03561 | 3599.77447 | 3198.63187 | 3572.94388 | 4566.91628 | 4558.58483 | 3085.93072 | 4938.98743 |
| tRNA-Thr-CGT | 4059.88984 | 1217.45574 | 3822.0167 | 3794.94297 | 3907.23968 | 6907.69151 | 1280.28321 | 701.532367 | 2987.8449 | 1644.57212 |
| tRNA-Gly-TCC | 2477.30858 | 3644.8054 | 1895.99482 | 2190.22423 | 2011.02103 | 3237.98039 | 3140.56049 | 2828.13833 | 3104.79338 | 3484.17363 |
| tRNA-Thr-AGT | 1634.2996 | 2574.80547 | 3826.30628 | 2153.04928 | 2339.59336 | 2687.15154 | 2895.6741 | 2363.20119 | 2603.0467 | 3716.10047 |
| tRNA-Trp-CCA | 2410.07473 | 3244.02804 | 1467.03671 | 2295.55326 | 2066.44287 | 2925.3478 | 2354.3463 | 2943.68483 | 3417.91349 | 3115.19911 |
| tRNA-Ser-CGA | 2363.52823 | 1886.67831 | 2299.21544 | 2298.65117 | 2343.55206 | 2322.41352 | 2371.53131 | 2305.42793 | 2508.73341 | 2756.76673 |
| tRNA-Ala-TGC | 2048.04634 | 1648.48045 | 1239.68892 | 1750.32063 | 1801.20978 | 2947.6787 | 3144.85674 | 2005.55724 | 2757.72049 | 2451.04499 |
| tRNA-Asn-GTT | 1396.39523 | 2053.03872 | 2127.63219 | 1939.2933 | 1840.79681 | 2344.74442 | 1971.97984 | 1719.44208 | 1829.67775 | 2498.48457 |
| tRNA-Met-CAT | 1649.81511 | 1739.22249 | 1411.27216 | 1790.5935 | 2050.60806 | 2277.75173 | 2221.16248 | 1942.28177 | 2127.70773 | 2509.0267 |
| tRNA-Leu-TAA | 1815.3138 | 1546.39565 | 1171.05562 | 1552.05423 | 1725.99443 | 2337.30079 | 2268.42126 | 2156.86814 | 2248.42874 | 2519.56883 |
| tRNA-Leu-CAA | 2079.07734 | 1550.17657 | 1038.07861 | 1316.61287 | 1706.20091 | 2419.18075 | 2401.60508 | 1614.9 | 2003.2142 | 2108.4258 |
| tRNA-Lys-TTT | 2601.4326 | 1493.46279 | 862.205788 | 1462.21476 | 1314.28933 | 2046.9991 | 1662.64967 | 1491.10017 | 2289.92659 | 1734.18022 |
| tRNA-Val-TAC | 677.510279 | 771.307366 | 403.220617 | 582.407574 | 522.548771 | 960.228668 | 992.4343 | 869.349913 | 1041.21868 | 1059.48396 |
| tRNA-Ile-AAT | 636.135605 | 415.901031 | 699.201709 | 535.938884 | 942.171269 | 930.454136 | 975.24929 | 577.732537 | 796.004135 | 911.894157 |
| tRNA-Thr-TGT | 610.276435 | 665.441649 | 639.147574 | 631.974176 | 665.062072 | 870.905071 | 915.101757 | 583.234752 | 780.914009 | 1054.2129 |
| tRNA-Phe-GAA | 387.887564 | 680.565323 | 441.826847 | 461.588981 | 403.787687 | 491.279784 | 330.811433 | 718.039011 | 486.656556 | 621.98561 |
| tRNA-Tyr-GTA | 217.217036 | 219.293271 | 145.845755 | 111.524855 | 245.439574 | 379.625288 | 433.92149 | 140.306473 | 294.257453 | 358.432385 |
| tRNA-Ile-TAT | 134.467689 | 113.427554 | 51.4749724 | 43.3707768 | 55.4218394 | 119.098129 | 150.368833 | 99.0398635 | 135.811132 | 247.740031 |
| tRNA-Ile-GAT | 15.5155026 | 18.9045923 | 25.7374862 | 27.8812136 | 63.339245 | 66.9926978 | 81.6287952 | 27.5110732 | 26.4077201 | 52.7106449 |
| tRNA-Tyr-ATA | 0 | 3.78091846 | 4.28958104 | 3.09791263 | 3.95870281 | 0 | 0 | 2.75110732 | 0 | 0 |

#### Table S4. tRNA isoacceptor expression [tpm].

| isoacceptor amino acid | HD1 | HD2 | HD3 | HD4 | HD5 | HD6 | HD7 | HD8 | HD9 | HD10 |
|---|---|---|---|---|---|---|---|---|---|---|
| Ala | 72457.397 | 86760.7359 | 77486.9918 | 77398.2491 | 54934.9189 | 75813.403 | 89546.7883 | 77581.2264 | 76559.7531 | 89244.3929 |
| Arg | 105401.981 | 109945.328 | 90994.8825 | 113851.387 | 127482.107 | 116716.167 | 121648.386 | 115298.908 | 124648.211 | 130590.623 |
| Asn | 1396.39523 | 2053.03872 | 2127.63219 | 1939.2933 | 1840.79681 | 2344.74442 | 1971.97984 | 1719.44208 | 1829.67775 | 2498.48457 |
| Asp | 80871.9712 | 60574.0947 | 110924.276 | 108786.3 | 105535.058 | 89517.1315 | 59357.0229 | 60562.8766 | 69142.9563 | 56300.2398 |
| Cys | 5120.11585 | 9879.53994 | 3543.19394 | 4594.20443 | 4675.22802 | 5158.43773 | 6715.04247 | 4993.25979 | 6730.1961 | 8918.64112 |
| Gln | 135077.965 | 94057.9085 | 128571.612 | 123727.532 | 118899.639 | 157425.396 | 97357.3752 | 78015.9014 | 105098.953 | 103344.49 |
| Gly | 85381.8107 | 141531.121 | 81030.1858 | 93959.69 | 78469.4071 | 76535.4354 | 98736.4722 | 114146.194 | 95169.6507 | 117486.756 |
| His | 4685.68178 | 8253.745 | 7142.15243 | 8274.52463 | 7082.11933 | 5835.80834 | 5847.19949 | 6522.87546 | 6526.4794 | 6309.4642 |
| Ile | 786.118797 | 548.233177 | 776.414168 | 607.190875 | 1060.93235 | 1116.54496 | 1207.24692 | 704.283474 | 958.222987 | 1212.34483 |
| Leu | 56926.3789 | 81709.4289 | 80043.5821 | 68978.1225 | 60861.097 | 61819.3728 | 73483.1007 | 81405.2656 | 60700.0309 | 87489.1284 |
| Lys | 57226.3453 | 38444.3789 | 42784.2813 | 53680.63 | 41788.0669 | 59683.0501 | 44607.9885 | 35244.4359 | 57074.6282 | 50428.274 |
| Met | 28165.809 | 36784.5557 | 26059.2048 | 33640.2332 | 26836.0464 | 47557.3718 | 39400.9306 | 44936.587 | 44847.8538 | 51825.1061 |
| Phe | 387.887564 | 680.565323 | 441.826847 | 461.588981 | 403.787687 | 491.279784 | 330.811433 | 718.039011 | 486.656556 | 621.98561 |
| Pro | 47425.7195 | 48002.5408 | 42857.2041 | 42230.7449 | 43320.0849 | 62139.449 | 45419.9802 | 32694.1594 | 39653.078 | 54892.8656 |
| SeC | 13932.9213 | 15724.8399 | 14790.4754 | 19185.3729 | 19239.2957 | 16160.1274 | 15161.4746 | 16223.2799 | 17900.6617 | 13525.5515 |
| Ser | 275839.777 | 223066.627 | 257404.889 | 213619.663 | 281800.26 | 186001.504 | 264713.59 | 290618.724 | 252846.375 | 184197.349 |
| Thr | 6304.46588 | 4457.70287 | 8287.47056 | 6579.96642 | 6911.89511 | 10465.7481 | 5091.05907 | 3647.96831 | 6371.80561 | 6414.88549 |
| Trp | 2410.07473 | 3244.02804 | 1467.03671 | 2295.55326 | 2066.44287 | 2925.3478 | 2354.3463 | 2943.68483 | 3417.91349 | 3115.19911 |
| Tyr | 217.217036 | 223.074189 | 150.135336 | 114.622767 | 249.398277 | 379.625288 | 433.92149 | 143.057581 | 294.257453 | 358.432385 |
| Val | 19983.9673 | 34058.5135 | 23116.5522 | 26075.1306 | 16543.4191 | 21914.0558 | 26615.2835 | 31879.8316 | 29742.6379 | 31225.786 |
